## Supplemental file for "Randomized Spatial PCA (RASP): a computationally efficient method for dimensionality reduction of high-resolution spatial transcriptomics data"

### RASP Supplemental Information

### List of Algorithms

1 Randomized SVD . . . . . . . . . . 3
2 KD-Tree Construction . . . . . . . . . . 4
3 Build distance matrix from KD-Tree . . . . . . . . . . 5
4 query_ball_point . . . . . . . . . . 5
5 local_density . . . . . . . . . . 6
6 CHAOS Function . . . . . . . . . . 7
7 1-Nearest Neighbor Function . . . . . . . . . . 7
8 Moran's I Calculation . . . . . . . . . . 8

### List of Tables

1 Datasets used for RASP evaluation . . . . . . . . . . 9
2 Parameter Selection . . . . . . . . . . 9

### 1 Supplemental Methods

#### 1.1 Efficient numerical methods and data structures utilized by RASP

##### Randomized PCA/SVD

The RASP algorithm depends on randomized principal component analysis (rPCA)[1] to perform computationally efficient dimensionality reduction on the gene expression data. For the **Python** version of RASP, this is implemented using the **sklearn** (version 1.5.2) package[2], which has an option for randomized PCA when calling the `PCA()` function. For the **R** version of RASP, this is implemented using the `rpca` function in the **rsvd** package (version 1.0.5) [1]. This randomized approach builds on the impressive results from the field of randomized numerical linear algebra (RNLA)[3] and allows an approximate reduced rank embedding to be computed for very large datasets at a computational cost that is orders-of-magnitude faster than can be achieved using non-stochastic truncated PCA implementations. Randomized PCA is implemented using a randomized singular value decomposition (rSVD) , which includes the following key steps (see Algorithm 1 for a detailed description):

- **Random sampling:** Generate a random projection matrix that reduces the dimensions of the original data while preserving its structure.
- **Compute a non-stochastic SVD on the random embedding:** Perform a standard deterministic SVD on this lower-dimensional representation of the data.
- **Generate an approximate SVD for the full matrix:** The results of the smaller SVD are used to generate an approximate truncated SVD for the original matrix.

##### K-dimensional tree (KD-Tree) operations

The RASP method also relies on the use of the KD-Tree data structure for organizing spatial coordinates in k-dimensional space for fast range or nearest neighbor searches. Construction of the KD-Tree structure is detailed in Algorithm 2. For the **Python** version of RASP, this algorithm is implemented using the `KDTree` function provided by the **Scipy** package [4] (version 1.14.1). For the **R** version of RASP, this algorithm is implemented using the `nn2` function from the **RANN** package (version 2.6.2).

The KD-Tree structure can be described at a high-level as follows:

- **Node Structure:** Each node $N$ contains:
  - A point $p \in \mathbf{R}^k$: representing the location in $k$-dimensional space.
  - Left child $N_{\text{left}}$ and right child $N_{\text{right}}$: representing subtrees corresponding to the divided space.

**Algorithm 1** Randomized SVD

Inputs:

- Matrix $A$ (size $m \times n$): the input data matrix.
- Integer $k$: the number of significant singular values to retain.
- Integer $p$: oversampling parameter (to improve accuracy).

Outputs:

- Matrix $U_k$ (size $m \times k$): left singular vectors.
- Matrix $S_k$ (size $k \times k$): diagonal matrix of singular values.
- Matrix $V_k$ (size $n \times k$): right singular vectors.

Notation:

- Matrix $\Omega$ (size $n \times r$): random projection matrix.
- Matrix $Y$ (size $m \times r$): projected matrix $A\Omega$.
- Matrices $Q$ and $R$: orthonormal basis and upper triangular matrix from QR decomposition.
- Matrix $B$ (size $r \times n$): matrix after projection onto $Q$.

$r = k + p$ ▷ Dimension of the random projection
Generate a random matrix $\Omega$ (size $n \times r$) ▷ Entries sampled from a Gaussian or uniform distribution over a specified range.
$Y = A \cdot \Omega$ ▷ Project original matrix
$[Q, R] \leftarrow \texttt{QR}(Y)$ ▷ QR decomposition
$B = Q^T \cdot A$ ▷ Form smaller matrix
$[U_b, S_b, V_b] \leftarrow \texttt{SVD}(B)$ ▷ Compute SVD of $B$
$U_k = Q \cdot U_b$ ▷ Form left singular vectors
$S_k = S_b[1 : k, 1 : k]$ ▷ Retain top $k$ singular values
$V_k = V_b[1 : k, :]$ ▷ Retain top $k$ right singular vectors
return $U_k, S_k, V_k$

- **Partitioning:** The space is partitioned based on the median point along the chosen dimension $d$ (where $d$ cycles through dimensions $1, 2, \ldots, k$):
  - If the current dimension is $d$, nodes are divided into left and right subtrees based on whether their points have coordinates less than or greater than the median along dimension $d$.
- **Recursive Construction:** This process continues recursively for each subtree until a specified stopping condition is met (e.g., a maximum number of points in a leaf node).

**Algorithm 2** KD-Tree Construction

Inputs:

- Set of points $P$: a list of points in $\mathbf{R}^k$
- Integer $depth$: current depth in the tree (or dimension to split on)

Outputs:

- Node $N$: root of the constructed KD-Tree

if size of $P$ is 0 then return None ▷ Base case: no points to construct a node
$k \leftarrow$ number of dimensions
$axis \leftarrow depth \mod k$ ▷ Choose axis for this level
Sort points $P$ along the chosen axis
$median \leftarrow \lfloor \texttt{size}(P)/2 \rfloor$ ▷ Find median index
$N \leftarrow$ new node $p_{median}$ ▷ Create a node at the median
$N.left \leftarrow \texttt{KD-Tree}(P[0 : median], depth + 1)$
$N.right \leftarrow \texttt{KD-Tree}(P[median + 1 : end], depth + 1)$
return $N$ ▷ Return the constructed node

The KD-Tree is used to calculate the sparse distance matrix leveraged for spatial smoothing. The KD-Tree can be queried to find distances to all neighbors within a specified distance, or find all distances to K-nearest-neighbors, see `Build distance matrix from KD-Tree`3.

The KD-Tree is also used when performing local cell density calculations. The **SciPy** function `query_ball_point()` is used to identify all points that fall within a specified radius r of a given point. The `nn2` function is used for this purpose in the **R** version of RASP. These functions exploit the KD-Tree's partitioning to quickly find neighboring points without needing to check every point in the dataset. For each point, the density is computed as the number of neighboring points found divided by the volume of the neighborhood. In a two-dimensional space, the volume is calculated using the area of a circle with radius r (i.e., $\pi r^2$). This formula ensures that the density reflects the spatial distribution of points around each queried point. See `local_density()`5 and `query_ball_point()`4.

#### 1.2 Adjusted Rand Index calculation

For the ovary, breast cancer, DLPFC, and simulated datasets we compared the identified spatial domains or cell type annotations directly to the ground truth labels using the adjusted rand index (ARI)[5] by using the `adjusted_rand_score` function from the **sklearn** package (version 1.5.2) and `adjustedRandIndex` from the **mclust** library (6.1.1) in the **R** version of RASP. Mathematically, the ARI is defined as:

$$\text{ARI} = \frac{\sum_{ij}\binom{n_{ij}}{2} - \left[\sum_i \binom{a_i}{2}\sum_j \binom{b_j}{2}\right]/\binom{n}{2}}{\frac{1}{2}\left[\sum_i \binom{a_i}{2} + \sum_j \binom{b_j}{2}\right] - \left[\sum_i \binom{a_i}{2}\sum_j \binom{b_j}{2}\right]/\binom{n}{2}}$$

Where:

- $n$ is the total number of elements (e.g., cells in spatial domains or annotations).
- $n_{ij}$ is the number of elements that are in cluster $i$ in the first partition and in cluster $j$ in the second partition.
- $a_i$ is the number of elements in cluster $i$ of the first partition.

**Algorithm 3** Build distance matrix from KD-Tree

**Inputs:**

- Set of points $P$: a list of points in $\mathbf{R}^k$
- Integer $k$: number of nearest neighbors to consider for KNN (if used, $k > 0$)
- Float $\epsilon$: maximum distance threshold (if used, $\epsilon > 0$)

**Outputs:**

- Sparse distance matrix $D$: a structure holding non-zero distances

$T \leftarrow$ KD-Tree($P$) ▷ Construct the KD-Tree from points
Initialize an empty sparse matrix $D$
**for** each point $p_i \in P$ **do**
    **if** $k > 0$ **then** ▷ Only kNN specified
        neighbors $\leftarrow$ kNN_search($T, p_i, k$)
    **else if** $\epsilon > 0$ **then** ▷ Only distance threshold specified
        neighbors $\leftarrow$ range_search($T, p_i, \epsilon$)
    **else**
        continue ▷ Skip this point if neither kNN nor threshold is specified
    **for** each neighbor $p_j$ in neighbors **do**
        $d \leftarrow$ Distance($p_i, p_j$) ▷ Calculate distance
        **if** $k > 0$ or $d < \epsilon$ **then**
            Add the distance $d$ to the sparse matrix $D$ at position $(i, j)$
**return** $D$ ▷ Return the sparse distance matrix

**Algorithm 4** query_ball_point

`Inputs:`

- *tree*: the constructed KD-Tree
- Point $p$: the center point for the search
- Float $r$: radius for searching neighbors

`Outputs:`

- List *indices*: indices of points within the radius $r$

Initialize an empty list *indices*
function QUERY_BALL_POINT(tree, p, r)
    neighbors $\leftarrow$ search_within_radius($tree, p, r$) ▷ Search KD-Tree for neighbors
    for each neighbor $n \in neighbors$ do
        if Distance($p, n$) < $r$ then
            Append the index of $n$ to *indices*
    return *indices*

**Algorithm 5** local_density

Inputs:

- Set of coordinates *coords*: a list of points in $\mathbf{R}^k$
- Float *neighborhood_size*: radius for density calculation

Outputs:

- List *densities*: density values for each point in *coords*

$tree \leftarrow$ KD-Tree($coords$) ▷ Construct the KD-Tree from coordinates
Initialize an empty list *densities*
**for each** point $p \in coords$ **do**
    $indices \leftarrow$ query_ball_point($tree, p, neighborhood\_size$) ▷ Find neighbors within radius
    $density \leftarrow \frac{\texttt{len}(indices)}{\pi \cdot (neighborhood\_size)^2}$ ▷ Calculate density
    $densities$ += $density$
**return** *densities* ▷ Return the list of densities

- $b_j$ is the number of elements in cluster $j$ of the second partition.
- $\binom{n}{2}$ is the binomial coefficient, representing the number of ways to choose 2 elements from $n$, calculated as:

$$\binom{n}{2} = \frac{n(n-1)}{2}$$

The ARI formula adjusts for the chance similarity between clusters by considering both pairwise agreements and disagreements. The numerator counts the agreements, and the denominator normalizes the score, yielding an index that ranges between -1 (no agreement) and 1 (perfect agreement), with 0 indicating random labeling.

#### 1.3 CHAOS score calculation

The spatial continuity and compactness of each clustering result is quantified by the CHAOS[6] score in place of the ARI. For the olfactory bulb dataset, the CHAOS score is used in place of ARI to access cluster quality and is detailed in Algorithm 6. The CHAOS score is based on the distances between points in a given cluster and their nearest neighbors, capturing how spatially compact the clusters are. The CHAOS score provides a measure of how spatially coherent the clusters are, with lower values indicating tighter and more compact clusters. The CHAOS score can be described at a high-level as follows:

- **Standardize Locations:** the location data is standardized such that each spatial coordinate has zero mean and unit variance.
- **1-Nearest Neighbor Calculation:** For each point a in cluster, we calculate the Euclidean distance to its nearest neighbor within the cluster. This is done by constructing a 1-Nearest Neighbor (1NN) graph. See Algorithm 7 for details.
- **Compute Cluster Distances:** For each cluster we sum the 1NN distances over all points in the cluster.
- **CHAOS Score Calculation:** The CHAOS score is then computed as the average 1NN distance across all clusters, weighted by the number of points in each cluster.

#### 1.4 Moran's I calculation

The spatial autocorrelation of each cluster is quantified using Moran's I[7], which assesses the degree of clustering of similar values across a spatial distribution, with positive values indicating clustering of similar values and negative values indicating dispersion. For the olfactory bulb dataset, Moran's I value is used in place of ARI to access cluster quality. The assignment of clusters is based on the proximity of spatial coordinates derived from k-nearest neighbors (kNN). The Moran's I statistic calculation is detailed in Algorithm 8

**Algorithm 6** CHAOS Function

Inputs:

- clusterlabel: Array of cluster labels for each data point.
- $L$: Array of spatial locations corresponding to each data point.

Outputs:

- CHAOS: The CHAOS score, representing the spatial compactness and continuity of the clusters.

NAs $\leftarrow$ indices where isna(clusterlabel) ▷ Identify indices of NA (null) values in cluster labels.
if length(NAs) $> 0$ then
    clusterlabel $\leftarrow$ delete(clusterlabel,NAs) ▷ Remove NA values from cluster label array.
    L $\leftarrow$ delete(L,NAs) ▷ Remove corresponding locations for deleted cluster labels.
L $\leftarrow$ scale(L) ▷ Standardize location data for better numerical stability.
unique_labels $\leftarrow$ unique(clusterlabel) ▷ Extract unique cluster labels from the cluster label array.
dist_val $\leftarrow$ zeros(length(unique_labels)) ▷ Initialize an array to hold distance values for each unique cluster.
for each cluster $k \in$ unique_labels do
    $L_k \leftarrow$ L[clusterlabel $==$ $k$] ▷ Extract locations corresponding to cluster $k$.
    if $L_k$ has only one point then
        Continue ▷ Skip cluster if it consists of only one point.
    for each index $i \in [0, \text{length}(L_k) - 1]$ do
        results[$i$] $\leftarrow$ fx_1NN($i, L_k$) ▷ Compute the nearest neighbor distance for point at index $i$.
    dist_val[$count$] $\leftarrow$ sum(results) ▷ Sum the 1NN distances for points in cluster $k$.
dist_val $\leftarrow$ dist_val[no NaN values] ▷ Remove any NaN values from the distance array.
Return CHAOS $\leftarrow \frac{\sum \text{dist\_val}}{\text{length(clusterlabel)}}$ ▷ Compute and return the average CHAOS score.

**Algorithm 7** 1-Nearest Neighbor Function

Inputs:

- $i$: Index of the point for which to compute the nearest neighbor.
- $L_k$: Array of spatial locations corresponding to points in the cluster.

Outputs:

- $d_{i,1}$: Euclidean distance from point at index $i$ to its nearest neighbor within the cluster.

distances $\leftarrow$ pairwise_distance([$L_k[i]$], $L_k$) ▷ Compute distance between the point at index $i$ and all points in $L_k$.
nearest_neighbor $\leftarrow$ copy(distances) ▷ Create a copy of the distances to sort them.
nearest_neighbor $\leftarrow$ partition(nearest_neighbor, 1) ▷ Partially sort distances such that the 1st nearest neighbor distance is in the correct position.
    return $d_{i,1} \leftarrow$ nearest_neighbor[0, 1] ▷ Return the distance to the nearest neighbor.

**Algorithm 8** Moran's I Calculation

`Inputs:`

- $C$: `Set of clusters in the dataset.`
- $L$: `Array of spatial coordinates corresponding to each data point in the dataset.`
- $Y$: `Vector of values assigned to each point based on their cluster.`
- $k$: `Number of nearest neighbors for kNN graph construction.`

`Outputs:`

- $I$: `Moran's I statistic, representing the degree of spatial autocorrelation.`

```
w ← kNN(L, k)   ▷ Construct kNN graph where w represents the weight matrix of the neighbors.
for each point i in the dataset do
    for each neighbor j in w do
        if j is one of the k nearest neighbors of i then
            w_ij ← 1                                  ▷ Assign weight of 1 for neighbor j.
        else
            w_ij ← 0                                  ▷ Assign weight of 0 if not a neighbor.
Ȳ ← (1/N) Σ_{i=1}^{N} Y_i                             ▷ Calculate the mean of the values in vector Y.
I ← N / (Σ_{i=1}^{N} Σ_{j=1}^{N} w_ij) · (Σ_{i=1}^{N} Σ_{j=1}^{N} w_ij (Y_i − Ȳ)(Y_j − Ȳ)) / (Σ_{i=1}^{N} (Y_i − Ȳ)^2)
                                                      ▷ Compute I using the provided formula.
    return I                                          ▷ Return the calculated Moran's I statistic.
```

$$\bar{Y} \leftarrow \frac{1}{N}\sum_{i=1}^{N} Y_i$$

$$I \leftarrow \frac{N}{\sum_{i=1}^{N}\sum_{j=1}^{N} w_{ij}} \cdot \frac{\sum_{i=1}^{N}\sum_{j=1}^{N} w_{ij}(Y_i-\bar{Y})(Y_j-\bar{Y})}{\sum_{i=1}^{N}(Y_i-\bar{Y})^2}$$

#### 1.5 Real world data preprocessing

**Mouse Ovary:** The dataset was provided directly by the authors [8] and was already preprocessed. RASP was applied directly to the `AnnData` object provided. Briefly, processing included cell segmentation (`CellPose` and `MERlin`)[9, 10] for the acquisition of cell and transcript data. Cells with fewer than 10 transcript counts were excluded from further analysis to ensure data quality. Data was log-normalized, scaled, PCA was performed, neighborhood identification, and cell clusterings calculated. To determine the cell type for each cluster, transcript counts were compiled for each gene across clusters, focusing on known markers within the top 10 identified transcripts. Differential expression analysis was conducted using **Seurat** v3 in **R**, allowing the identification of specific markers associated with each Leiden cluster. Additionally, visualization of spatial regions was achieved using **Squidpy** [11] in conjunction with the **AnnData** [12] and **Scanpy** [13] libraries.

**DLPFC:** We downloaded the raw DLPFC data (slide #151673) from the spatialLIBD website (`http://research.libd.org/spatialLIBD/index.html`). The dataset was loaded into **R** using **Seurat's** `CreateSeuratObject` function with the `min.cells = 20` and `min.features = 20` parameters. Data were processed using the `SCTransform` pipeline with the following parameters: `variable.features.n = NULL, variable.features.rv.th = 1.3,return.only = FALSE`. Data were then converted to the h5ad file format using **SeuratDisk** package's `SaveH5Seurat` and `Convert` functions to be processed via RASP in **Python**.

**Mouse Olfactory bulb:** The dataset was downloaded from the SEDR publication GitHub repository [14] (`https://github.com/JinmiaoChenLab/SEDR_analyses/tree/master/data`). Data were processed in **Python** using **Scanpy** unless otherwise specified. The standard processing pipeline was run, including the following functions: `calculate_qc_metrics`, `filter_genes` with `min_cells` = 50, `highly_variable_genes` with `flavor` = 'seurat', `normalize_total`, and `log1p`. The processed data were then analyzed using RASP.

**Breast Cancer:** We downloaded the Breast cancer Xenium dataset from the **SubcellularSpatialData R** package via **ExperimentHub**, dataset # EH8567, sample ID 'IDC' (`https://www.bioconductor.org/packages/release/data/experiment/html/SubcellularSpatialData.html`). Transcripts at each subcellular location were assigned to cells using the `tx2spe` function with `bin = 'cell'`. The resulting `SingleCellExperiment` object was converted to a *Seurat* object, and then converted to an .h5ad file using **SeuratDisk** package's `SaveH5Seurat` and `Convert` functions. Downstream analysis was done in **Python** using the **Scanpy** and **Squidpy** packages. Mitochondrial, ribosomal, and hemoglobin genes identified with the `var_names.str.startswith("MT-"),var_names.str.star` commands, quality control metrics where calculated with the `calculate_qc_metrics` command, and data were filtered using the `filter_cells` command where `min_genes = 20` and `min_cells = 3`. The filtered data were then log normalized and the top 3000 most highly variable genes where identified using the `highly_variable_genes` function with *favor = 'seurat'*. The processed data were then analyzed using RASP.

**Sagittal mouse brain:** We downloaded the sagittal mouse brian dataset from the Allen Institute spatial brain atlas API via the **abc_atlas_access Python** package. Specifically we followed the *MERFISH whole mouse brain spatial transcriptomics (Xiaowei Zhuang)* tutorial `https://alleninstitute.github.io/abc_atlas_access/notebooks/` `zhuang_merfish_tutorial.html` to access section 3.010 from the 'Zhuang-ABCA-3' dataset. This tutorial resulted in the raw data, along with the spatial annotations associated with each observation. Downstream analysis was done in **Python** using the **Scanpy** and **Squidpy** packages. quality control metrics where calculated with the `calculate_qc_metrics` command, and data were filtered using the `filter_cells` command where `min_counts = 10` and `filter_genes` where `min_cells=5`. The filtered data were then log normalized. The processed data were then analyzed using RASP.

Table 1: Datasets used for RASP evaluation

| **Platform** | **Tissue** | **Organism** | **n cell/n location** | **n genes** | **Ground truth annotation?** |
|---|---|---|---|---|---|
| MERFISH | Ovary | Mouse | 43,038 | 228 | True |
| Stereo-Seq | Brain (Olfactory bulb) | Mouse | 19,109 | 14,367 | False |
| 10x Xenium | Breast cancer | Human | 565,916 | 541 | True |
| 10x Visium | Brain (DLPFC) | Human | 3,638 | 15,124 | True |
| SRTsim | Simulation 1 | NaN | 10,000 | 150 | True |
| SRTsim | Simulation 2 | NaN | 10,000 | 150 | True |

##### 1.6 Simulated dataset processing

Raw simulated count data were processed in **Python** using the **Scanpy** and **Squidpy** packages. Count matrices were normalized to medial total counts using the `pp.normalize_total` command, logarithmized with the `pp.log1p` command and variable genes identified using the `pp.highly_variable_genes` function, all with default parameters. The processed data were then analyzed using RASP. Note: for testing the **SpatialPCA** algorithm on the simulated data, the raw simulated count matrix was processed via **Seurat's** `SCTransform` pipeline with default parameters, as per the packages recommendation.

Table 2: Parameter Selection

| **Parameter Name** | **Algorithm** | **Type** | **Range** | **Default** | **Description** | **Recommendations** |
|---|---|---|---|---|---|---|
| n_components | rSVD | int | 5-100 | 20 | Number of components to keep. | For spatial domain detection, use a range of $5-20$ PCs to capture larger domains. For clustering tasks with higher resolution or heterogeneity, users may want to increase this parameter to a range of $30-60$. Note that increased components will increase runtime. |

*Continued on next page*

| Parameter Name | Algorithm | Type | Range | Default | Description | Recommendations |
|---|---|---|---|---|---|---|
| n_oversamples | rSVD | int | 2-10 | 10 | Corresponds to the additional number of random vectors to sample the range of X to ensure proper conditioning. | Default is recommended for most situations. Smaller number can improve speed but negatively impact approximation quality. Users might want to increase this parameter up to $2k - n_{\text{components}}$ where $k$ is the effective rank, for large matrices, noisy problems, matrices with slowly decaying spectrum, or to increase precision accuracy. |
| power_ iteration_ normalizer | rSVD | string | auto, QR, LU, none | auto | Power iteration normalizer for randomized SVD solver. | Default is recommended. |
| threshold | RASP | int | 1-200 | 10 | Corresponds to the kNN threshold used to calculate the sparse distance matrix used for spatial smoothing. | For larger spatial domains and tissue sections, a larger smoothing threshold between 50 and 100 is recommended. For high-resolution cell type tasks, kNNs between 3 and 10 are recommended. Users should incorporate a priori knowledge of tissue structure. |
| n_neighbors | RASP | int | 5-50 | 10 | Number of kNN when constructing a NN graph of the smoothed principal components, used by the clustering algorithms. | Default of 10 is recommended. |
| n_clusters | RASP | int | inf | 10 | The number of clusters identified by the clustering algorithm. | - |

*Continued on next page*

| Parameter Name | Algorithm | Type | Range | Default | Description | Recommendations |
|---|---|---|---|---|---|---|
| covariates | RASP | string, list | nan | None | Specification of additional covariates, should be in the adata.obs slot of the adata object. | Users can include additional features to incorporate into the reduction and clustering step such as the number of counts (library size), cellular density, chromatin accessibility, or protein abundance measures. Incorporation of these features should be tested against the RNA alone and results compared with CHAOS score and Moran's I. |
| covariate_kNN | RASP | int, list | 1-200 | None | If covariates are included, covariate smoothing is the kNN threshold to use for smoothing the additional feature(s), or threshold to calculate local density. This can be the same or different than the smoothing threshold feature. | If users are adding local cell density as a covariate, then a different smoothing threshold is recommended than that used for the RNA smoothing. For other covariates, keeping the same smoothing threshold as the RNA is recommended. |

*Continued on next page*

| Parameter Name | Algorithm | Type | Range | Default | Description | Recommendations |
|---|---|---|---|---|---|---|
| cluster_ algorithm | Clustering | string | walktrap, mclust, louvain, leiden | louvain | Clustering algorithm utilized by RASP. | Clustering is largely dependent on the tissue type and structure users are hoping to identify. The walktrap algorithm works well for large, homogeneous structures, and evenly spaced spots such as 10x Visium, but has a large computational cost that increases runtime with larger datasets. Mclust utilizes GMM and generally performs consistently across all tissue types but is sensitive to the model type being fit. See the 'model_type' parameter. Louvain and Leiden algorithms outperform the other methods for high-resolution or heterogeneous labeling tasks such as identifying cell types or small interspersed tissue domains. |
| ground_ truth_ labels | Clustering | string | nan | None | Name of the ground truth labels in the adata.obs slot. | If the dataset is annotated, users can specify the data slot of the annotations and test the accuracy of RASP predictions. Used for benchmarking. |
| model_type | Mclust | string | See mclust-Model-Names for available model types | EEE | Corresponds to the GMM used by mclust. | Generally, EEE has the best performance on ST data. |

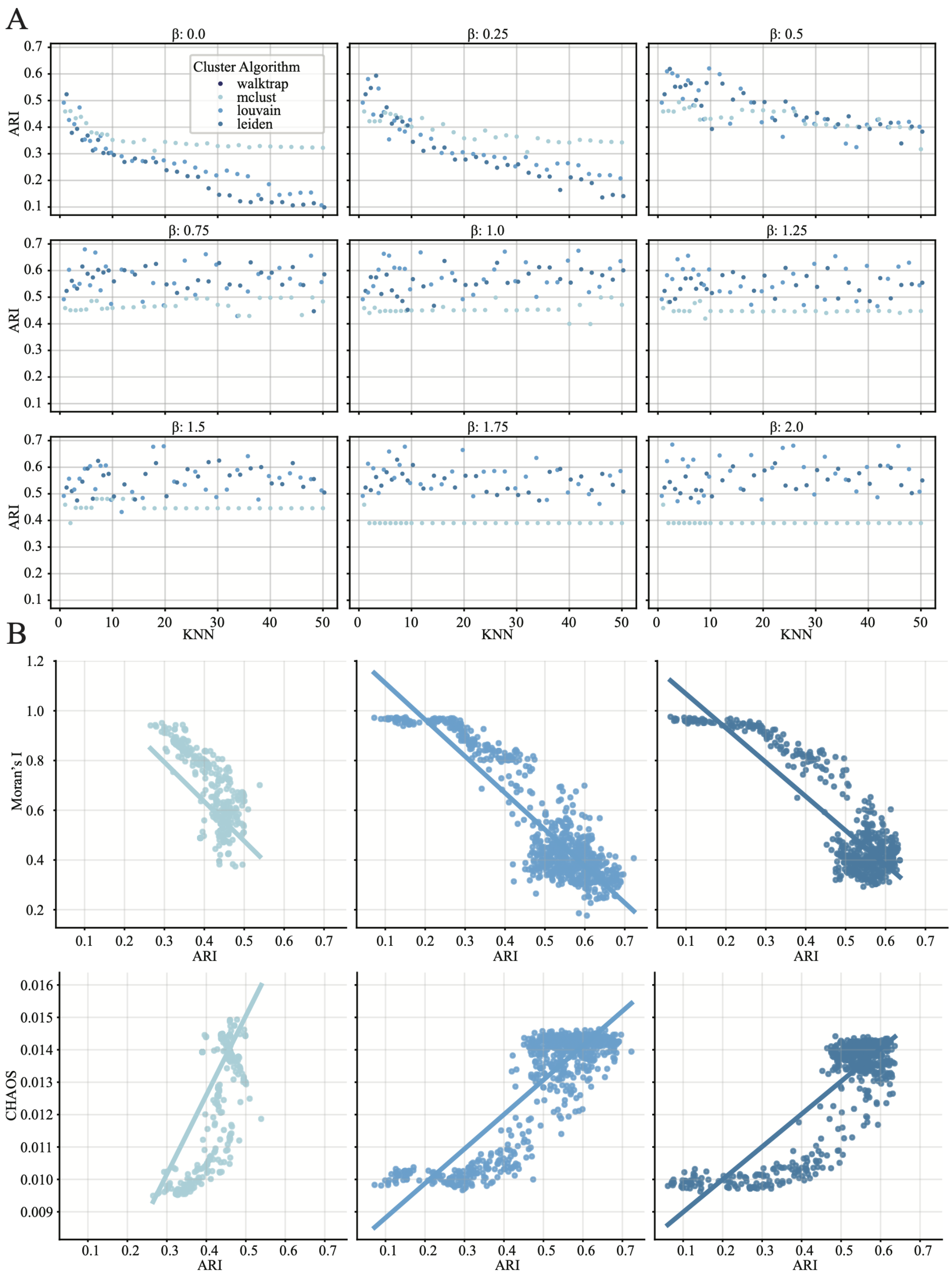

Extended Data Fig. 1: **Mouse Ovary supplement 1.** **A:** ARI values plotted against kNN distance threshold. Colors indicate clustering algorithm, each subplot corresponds to a distinct $\beta$ value. **B:** Moran's I and CHAOS values plotted against ARI value, colors indicate clustering algorithm. Lines represent robust linear regression best fit.

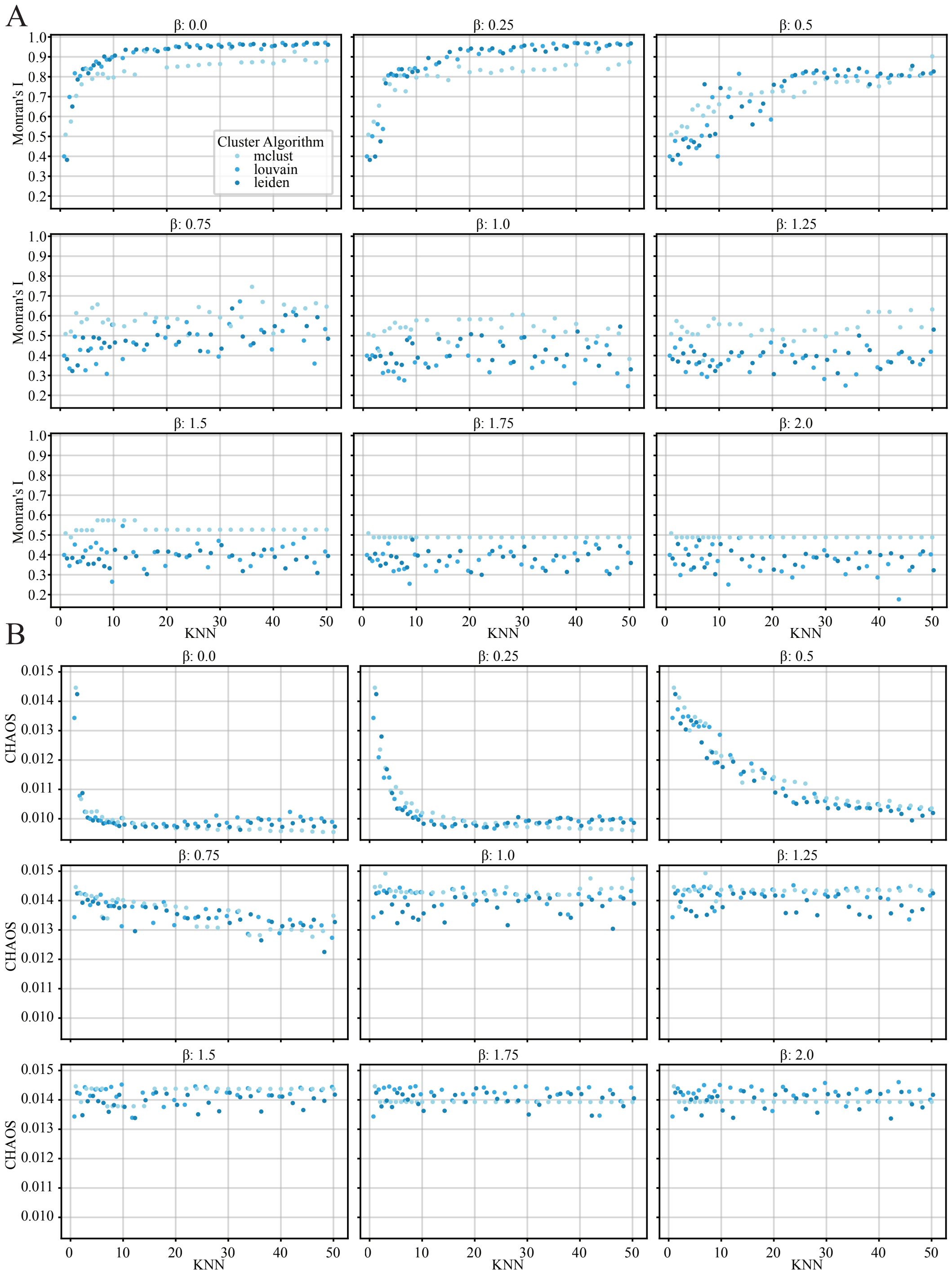

Extended Data Fig. 2: **Mouse Ovary supplement 2. A:** Moran's I statistic plotted against kNN distance threshold. Colors indicate clustering algorithm, each subplot corresponds to a distinct $\beta$ value. **B:** CHAOS score plotted against kNN distance threshold. Colors indicate clustering algorithm, each subplot corresponds to a distinct $\beta$ value.

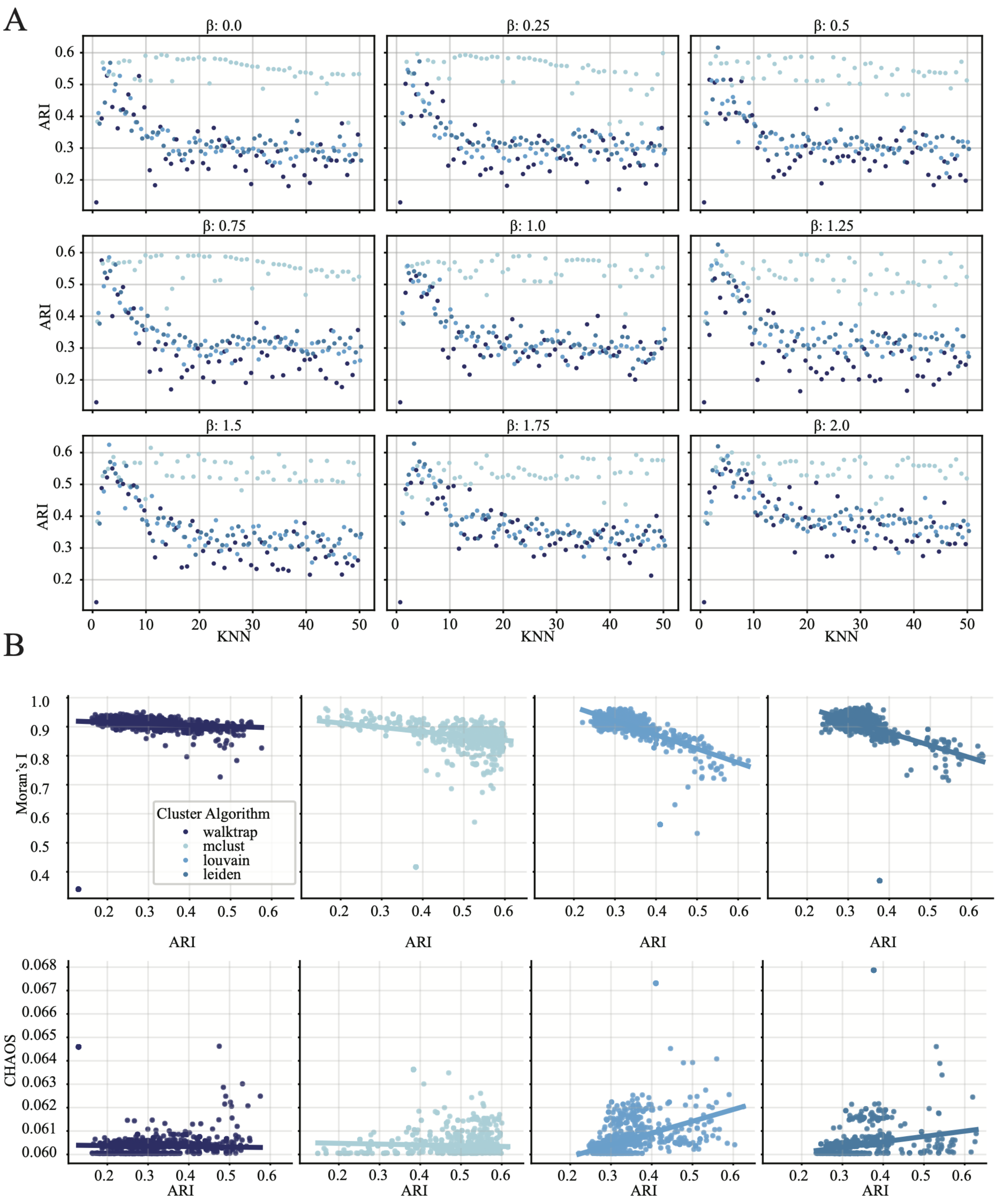

Extended Data Fig. 3: **DLPFC supplement 1. A:** ARI values plotted against kNN distance threshold. Colors indicate clustering algorithm, each subplot corresponds to a distinct $\beta$ value. **B:** Moran's I and CHAOS values plotted against ARI value, colors indicate clustering algorithm. Lines represent robust linear regression best fit.

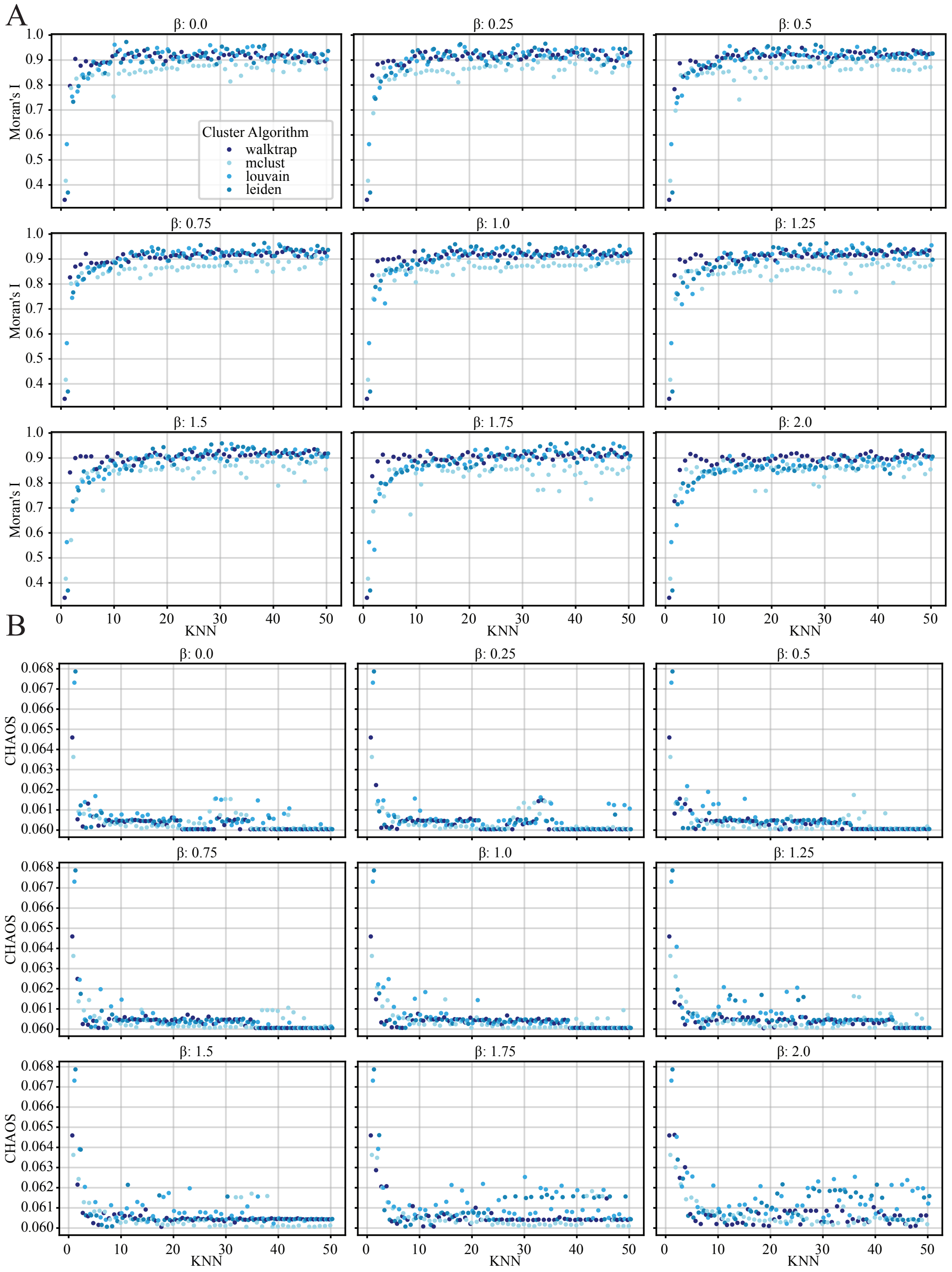

Extended Data Fig. 4: **DLPFC supplement 2. A:** Moran's I statistic plotted against kNN distance threshold. Colors indicate clustering algorithm, each subplot corresponds to a distinct $\beta$ value. **B:** CHAOS score plotted against kNN distance threshold. Colors indicate clustering algorithm, each subplot corresponds to a distinct $\beta$ value.

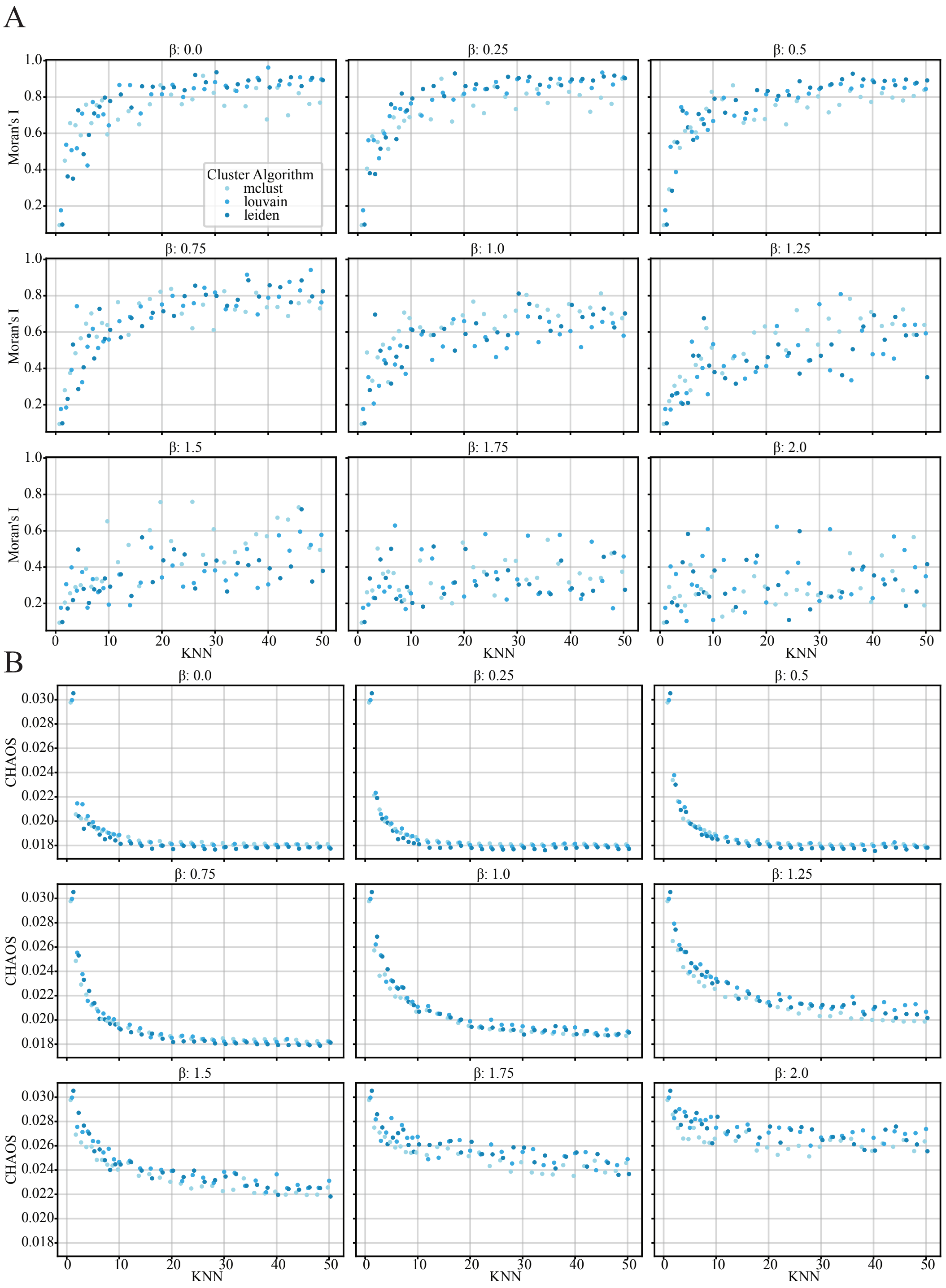

Extended Data Fig. 5: **Olfactory bulb supplement.** **A:** Moran's I statistic plotted against kNN distance threshold. Colors indicate clustering algorithm, each subplot corresponds to a distinct $\beta$ value. **B:** CHAOS score plotted against kNN distance threshold. Colors indicate clustering algorithm, each subplot corresponds to a distinct $\beta$ value.

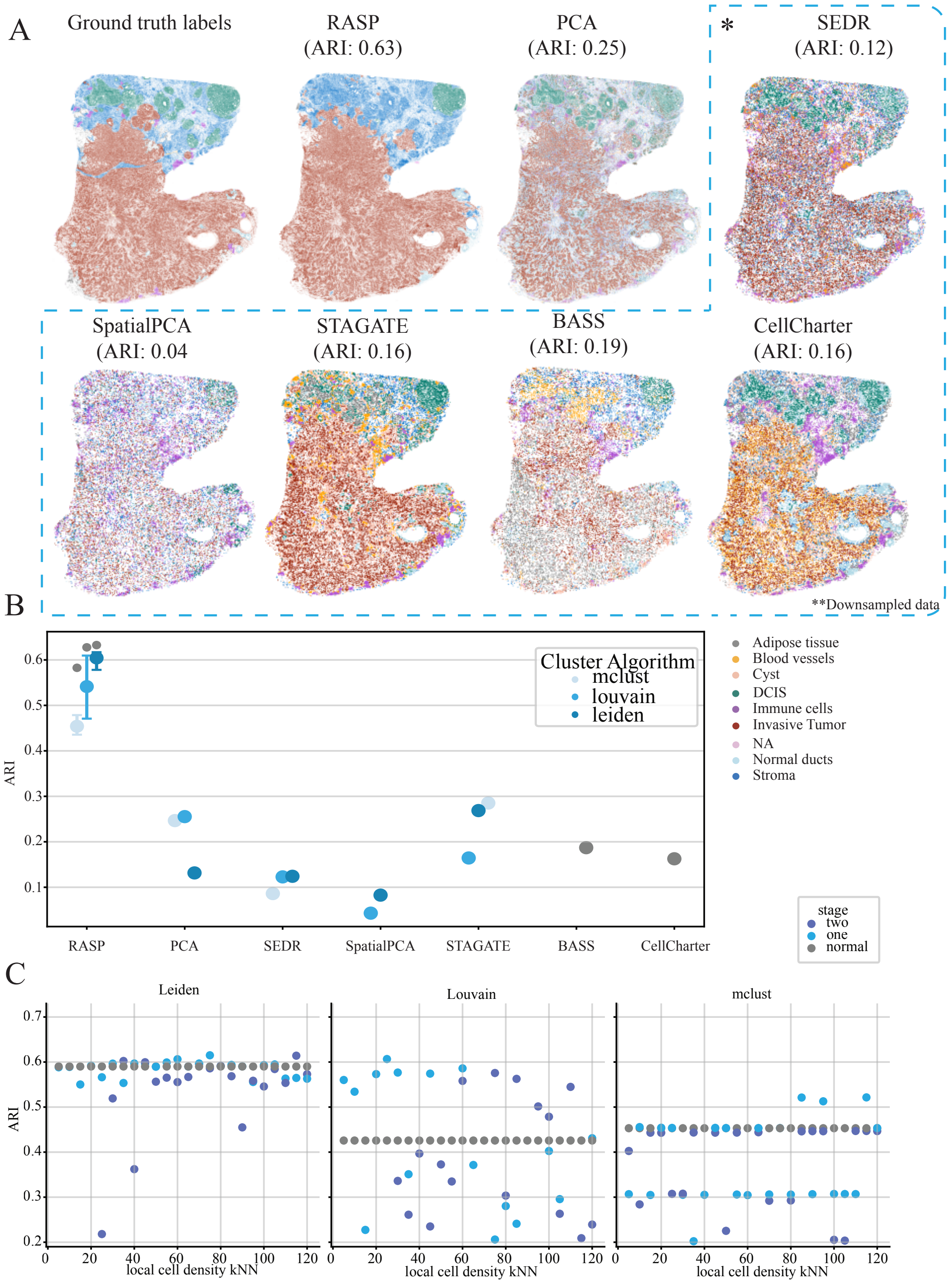

Extended Data Fig. 6: **Xenium supplement 1. A:** Ground truth annotations (left) and corresponding spatial domains identified by RASP, PCA, and other methods. Sections in the dotted line have been sketched to 10% of the original dataset. **B:** Quantification of ARI for all methods, different colors indicate the clustering algorithm used to assign labels. Interquartile range and median ARI values at default RASP parameters (kNN =60-100, $\beta$ = 0) is shown, along with maximum ARI values achieved by RASP. **C:** Quantification of ARI with addition of covariates for Leiden, Louvain and Mclust clustering algorithms. Colors indicate no covariates added (grey), one stage (blue), and two stage (purple) versions of RASP.

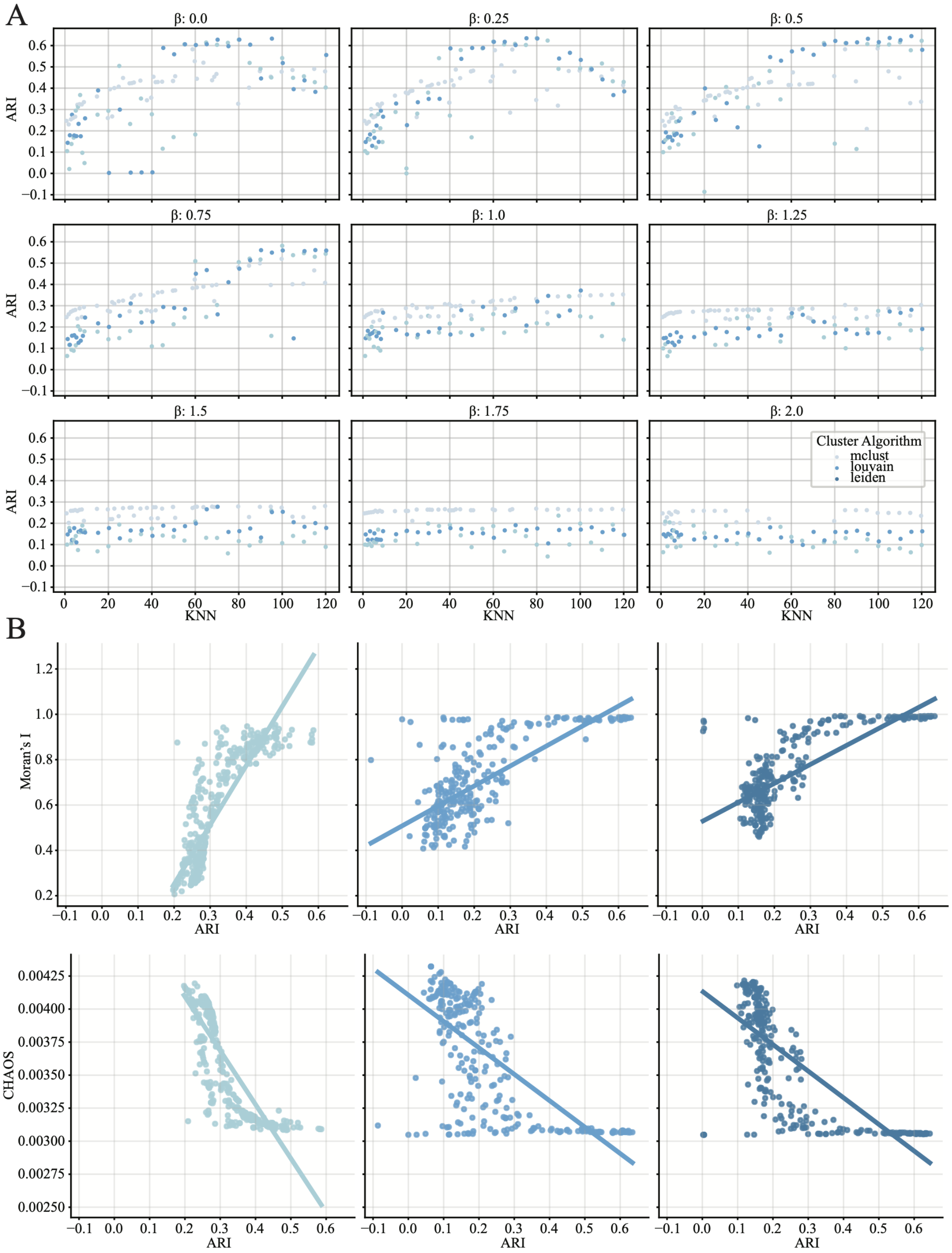

Extended Data Fig. 7: **Xenium supplement 2. A:** ARI values plotted against kNN distance threshold. Colors indicate clustering algorithm, each subplot corresponds to a distinct $\beta$ value. **B:** Moran's I and CHAOS values plotted against ARI value, colors indicate clustering algorithm.

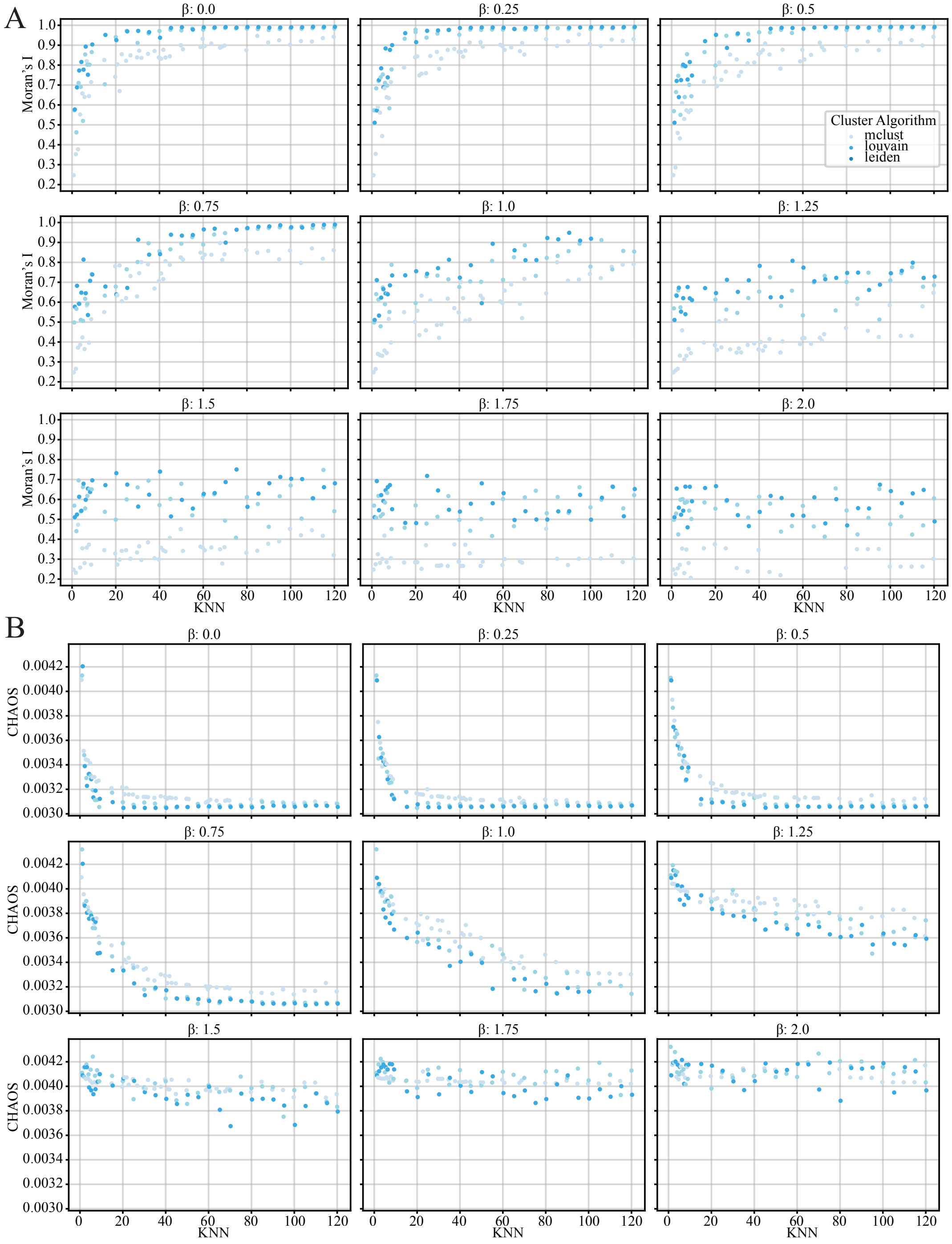

Extended Data Fig. 8: **Xenium supplement 3. A:** Moran's I statistic plotted against kNN distance threshold. Colors indicate clustering algorithm, each subplot corresponds to a distinct $\beta$ value. **B:** CHAOS score plotted against kNN distance threshold. Colors indicate clustering algorithm, each subplot corresponds to a distinct $\beta$ value. Lines represent robust linear regression best fit.

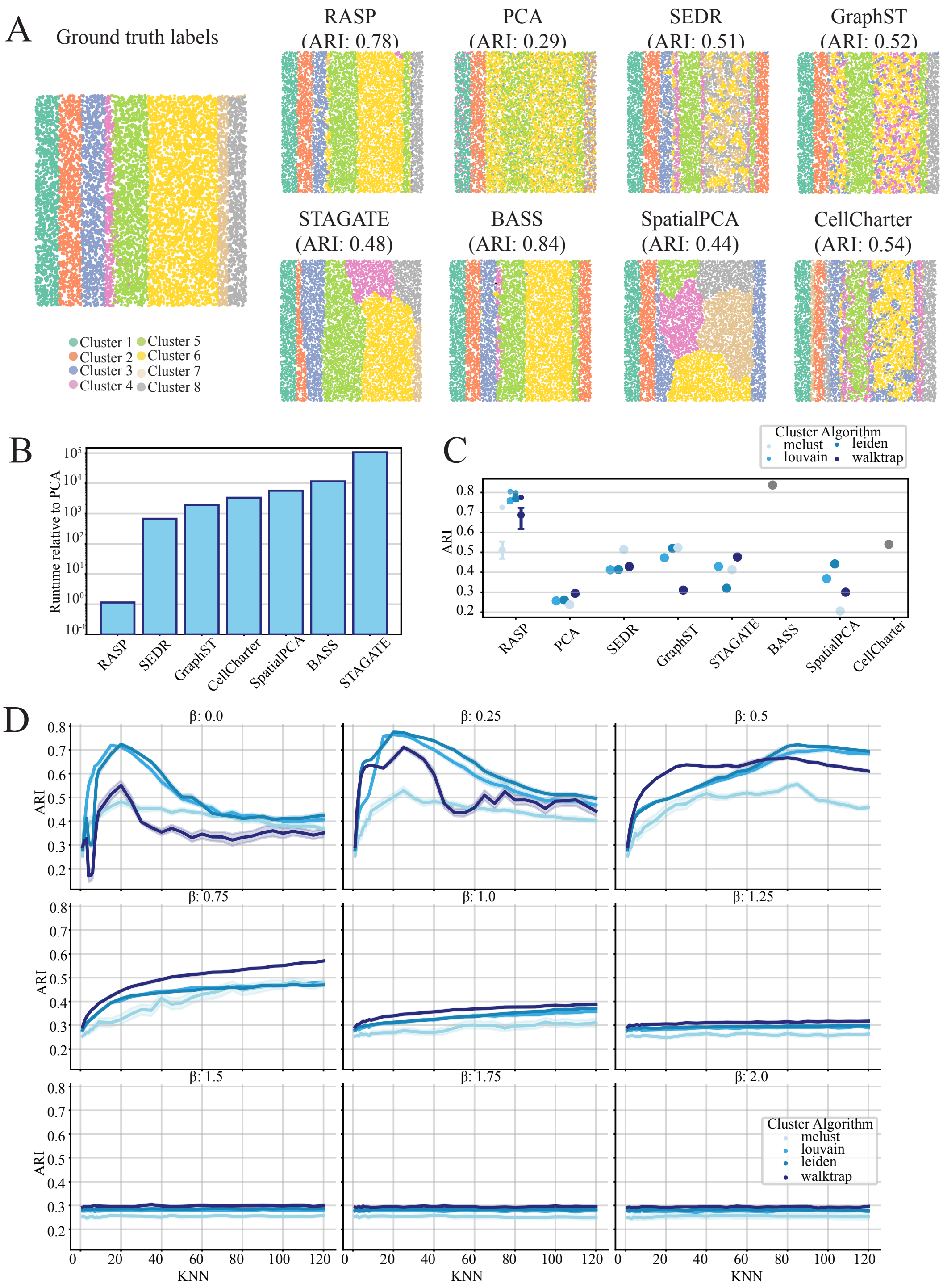

Extended Data Fig. 9: **Simulation 1 (Stripes) A:** Ground truth annotations (left) and corresponding spatial domains identified by RASP, PCA, and other methods. **B:** Quantification of runtime for all methods compared to normal PCA. **C:** Quantification of ARI for all methods, different colors indicate the clustering algorithm used to assign labels. Interquartile range and median ARI values at default RASP parameters (kNN =15-30, $\beta = 0.25$) is shown, along with maximum ARI values achieved by RASP. **D:** ARI values for RASP evaluated on 100 *Stripe* replicates plotted against kNN, each subplot represents a distinct $\beta$ value. Colored line represents the median, shaded colors indicate the interquartile range. Colors correspond to clustering algorithm.

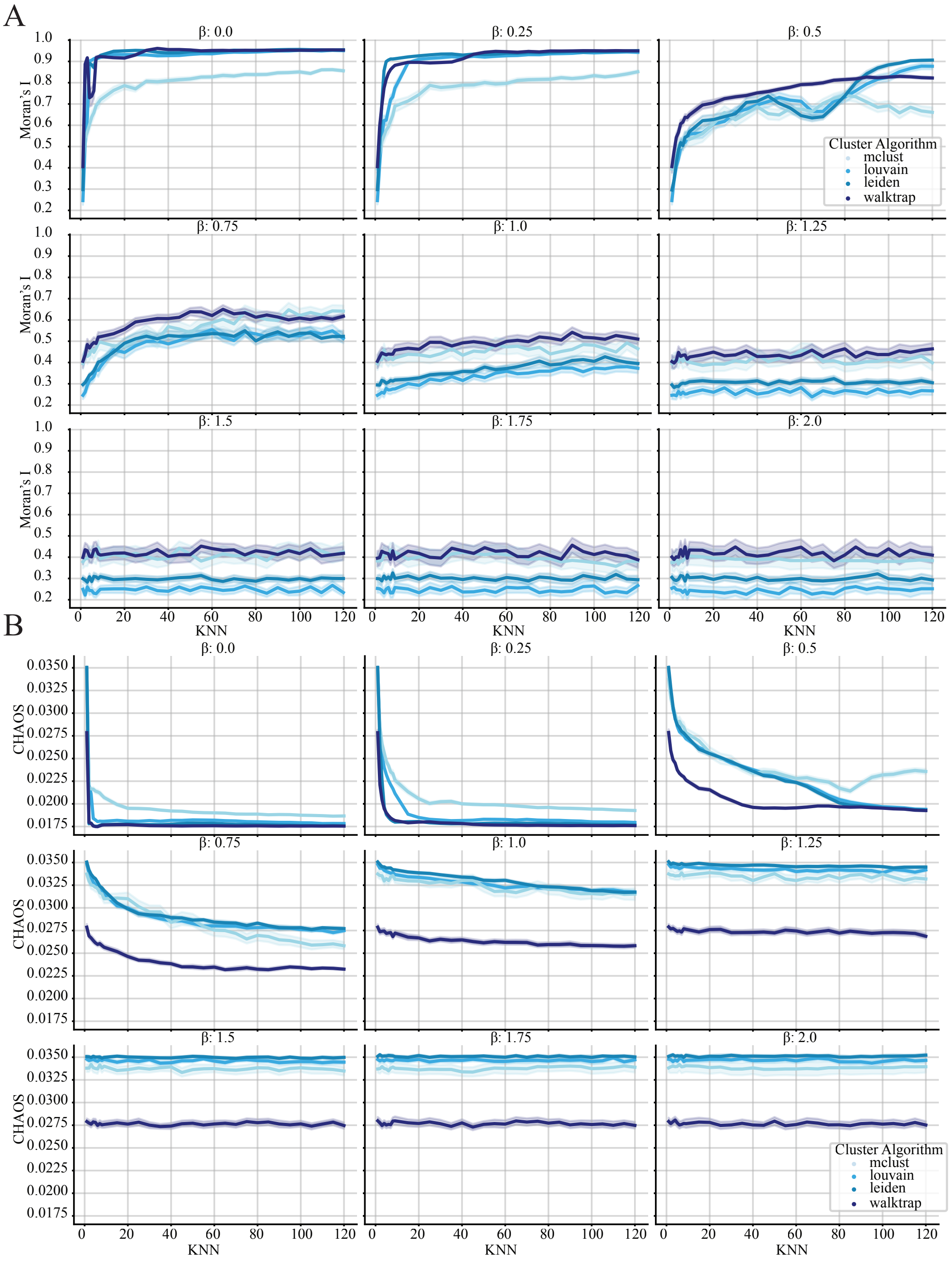

Extended Data Fig. 10: **Simulation 1 (Stripes)** **A:** Moran's I statistic plotted against kNN distance threshold. Colors indicate clustering algorithm, each subplot corresponds to a distinct $\beta$ value. **B:** CHAOS score plotted against kNN distance threshold. Colored line represents the median, shaded colors indicate the interquartile range. Colors correspond to clustering algorithm, each subplot corresponds to a distinct $\beta$ value.

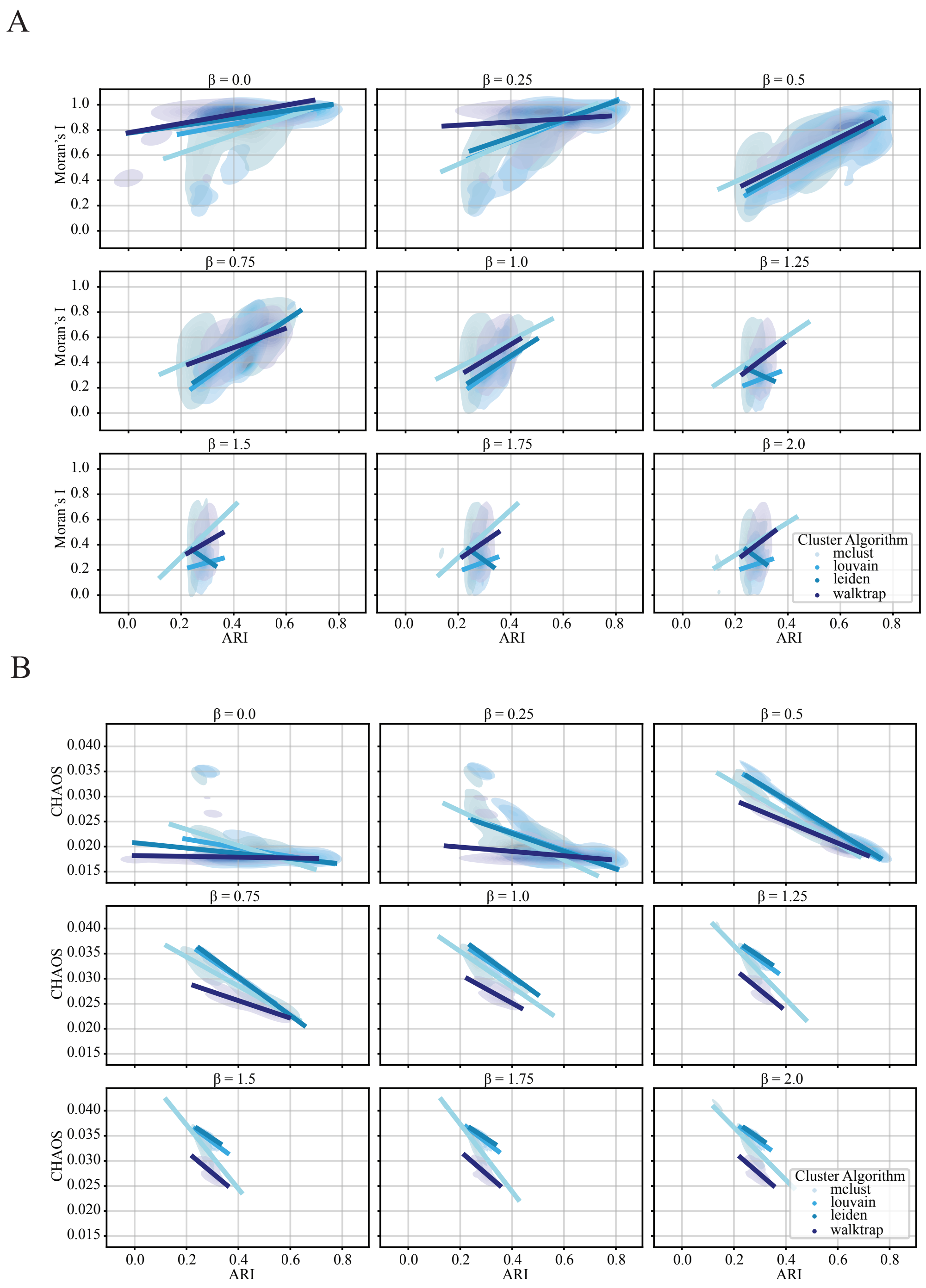

Extended Data Fig. 11: **Simulation 1 (Stripes) A:** Bivariate kernel density estimate of Moran's I statistic against ARI. Colors indicate clustering algorithm, each subplot corresponds to a distinct $\beta$ value. **B:** Bivariate kernel density estiamte of CHAOS score plotted against ARI. Colors indicate clustering algorithm, each subplot corresponds to a distinct $\beta$ value. Lines represent robust linear regression best fit.

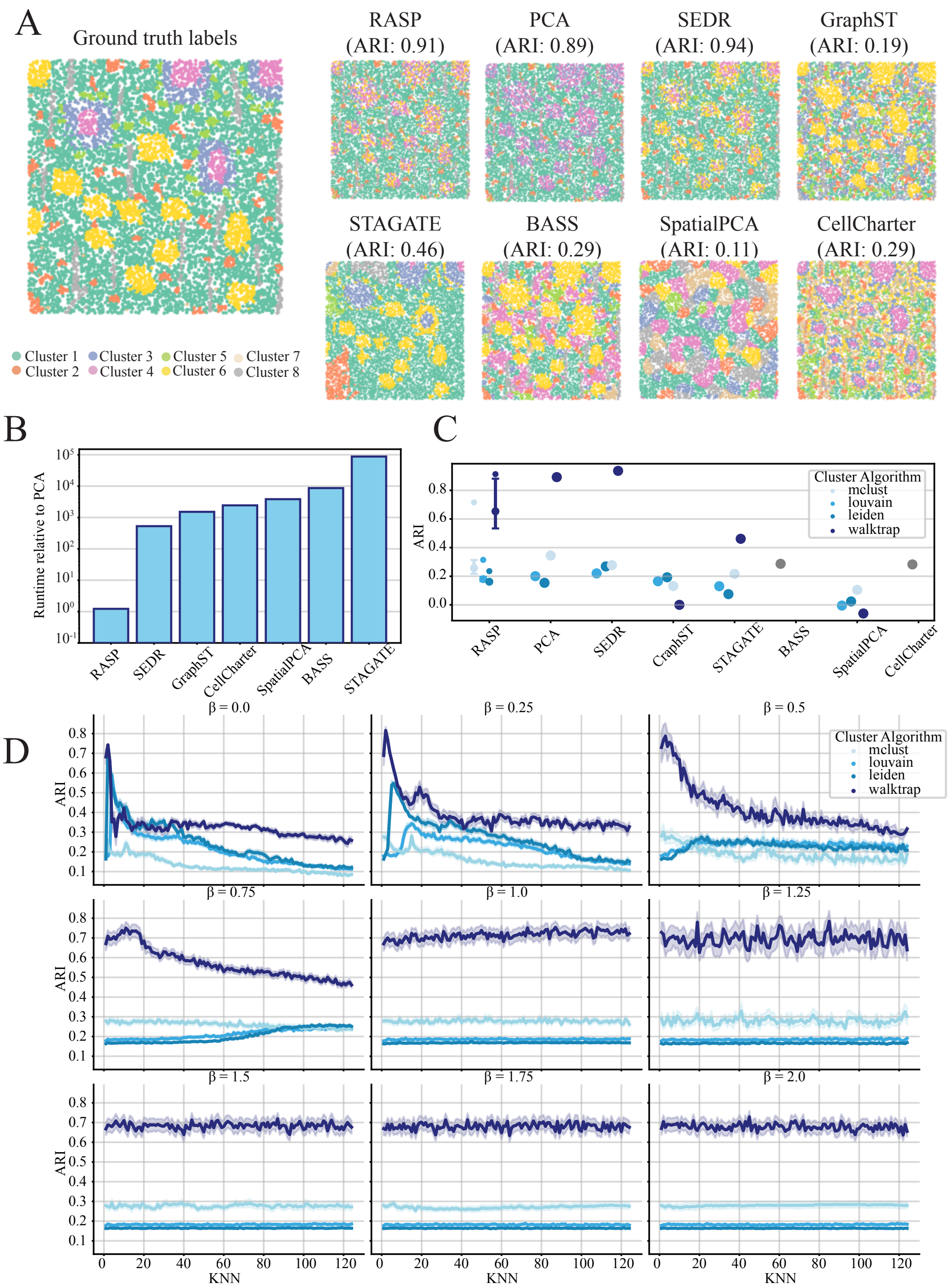

Extended Data Fig. 12: **Simulation 2 (Dots) A:** Ground truth annotations (left) and corresponding spatial domains identified by RASP, PCA, and other methods. **B:** Quantification of runtime for all methods compared to normal PCA. **C:** Quantification of ARI for all methods, different colors indicate the clustering algorithm used to assign labels. Interquartile range and median ARI values at default RASP parameters (kNN =3-20, $\beta$ = 2) is shown, along with maximum ARI values achieved by RASP. **D:** ARI values for RASP evaluated on 100 *Dot* replicates plotted against kNN, each subplot represents a distinct $\beta$ value. Colors indicate clustering algorithm.

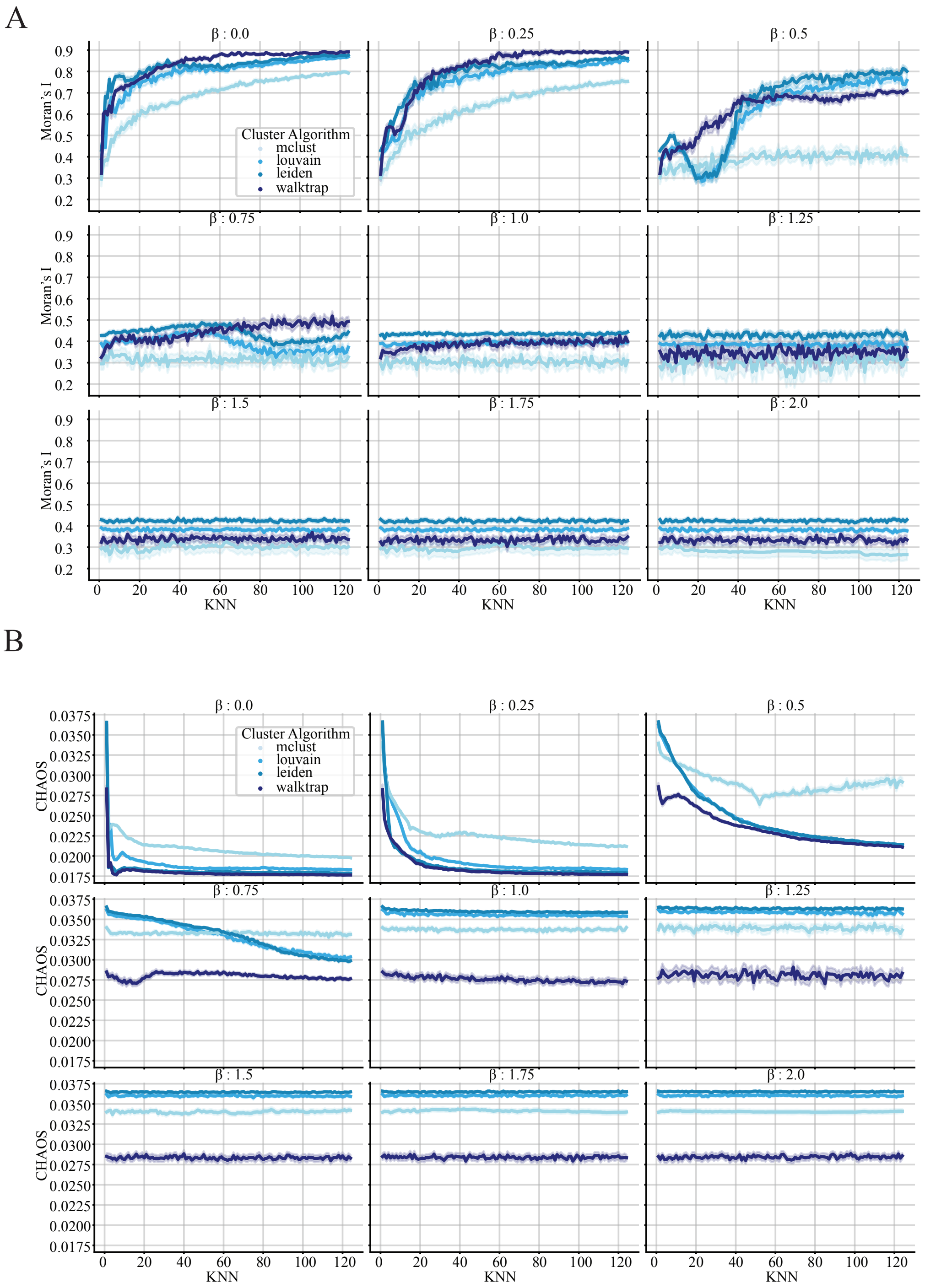

Extended Data Fig. 13: **Simulation 2 (Dots) A:** Moran's I statistic plotted against kNN distance threshold. Colors indicate clustering algorithm, each subplot corresponds to a distinct $\beta$ value. **B:** CHAOS score plotted against kNN distance threshold. Colored line represents the median, shaded colors indicate the interquartile range. Colors correspond to clustering algorithm, each subplot corresponds to a distinct $\beta$ value.

A

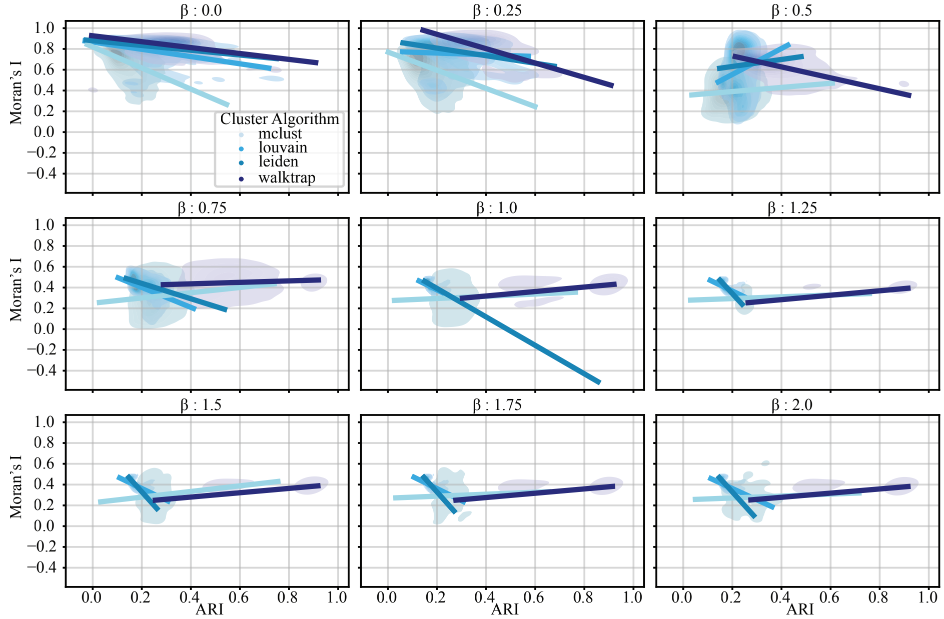

B

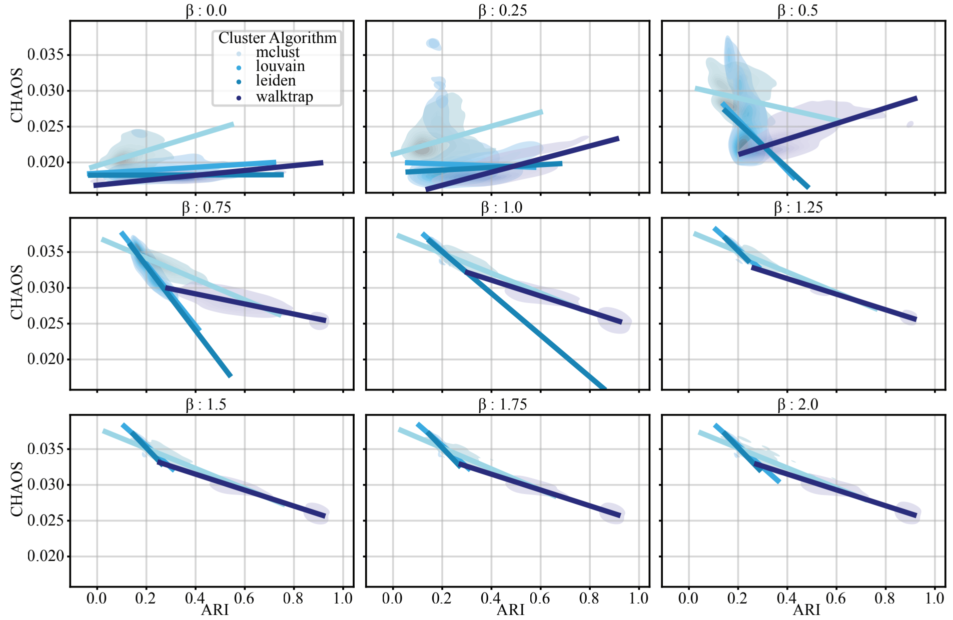

Extended Data Fig. 14: **Simulation 2 (Dots) A:** Bivariate kernel density estimate of Moran's I statistic against ARI. Colors indicate clustering algorithm, each subplot corresponds to a distinct $\beta$ value. **B:** Bivariate kernel density estiamte of CHAOS score plotted against ARI. Colors indicate clustering algorithm, each subplot corresponds to a distinct $\beta$ value. Lines represent robust linear regression best fit.

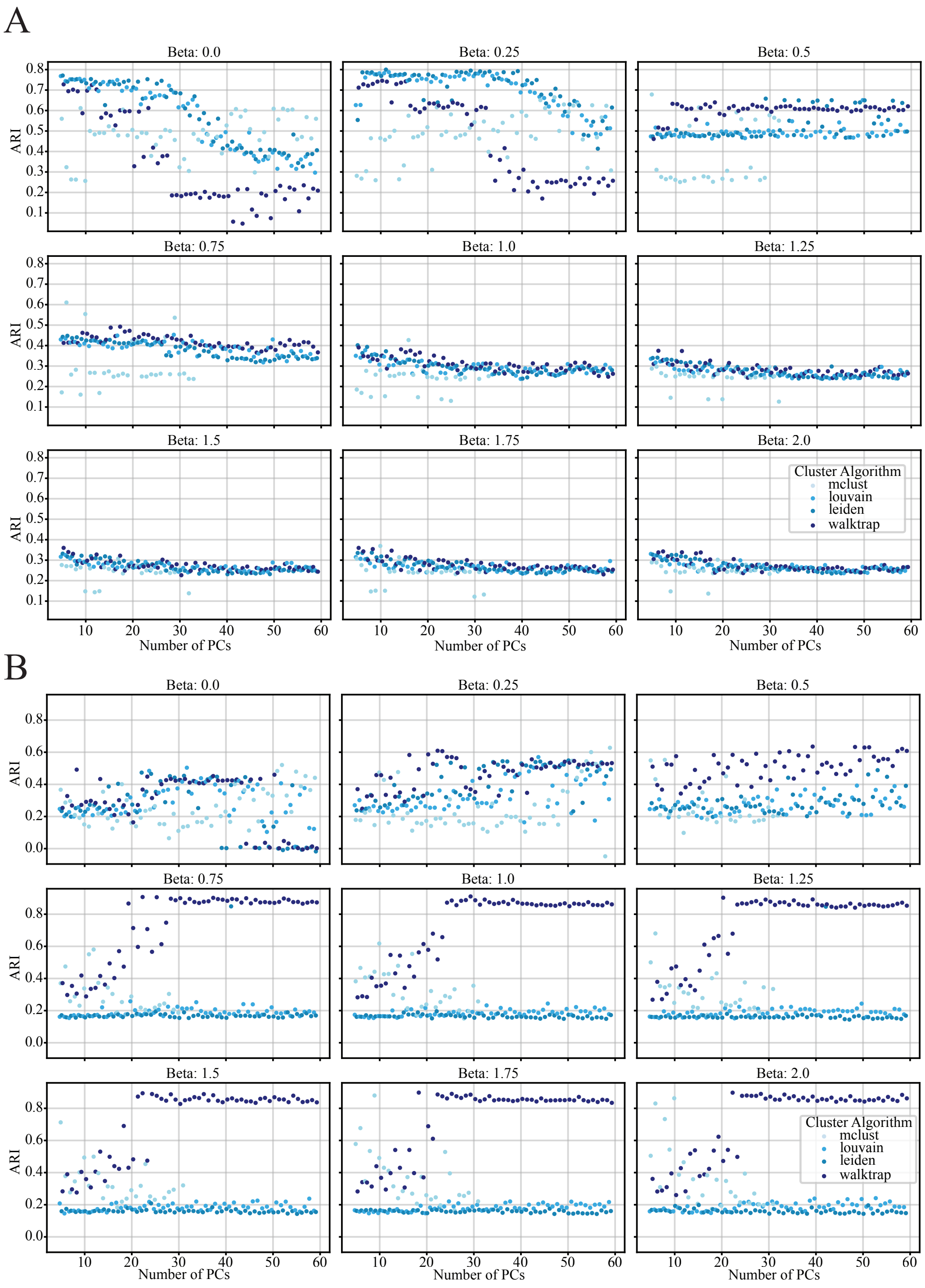

Extended Data Fig. 15: **PC sensitivity (simulated data) A:** ARI values plotted against number of PCs utilized by RASP for the *Stripes* dataset. Colors indicate clustering algorithm, each subplot corresponds to a distinct $\beta$ value. **B:** ARI values plotted against number of PCs utilized by RASP for the *Dots* dataset. Colors indicate clustering algorithm, each subplot corresponds to a distinct $\beta$ value.

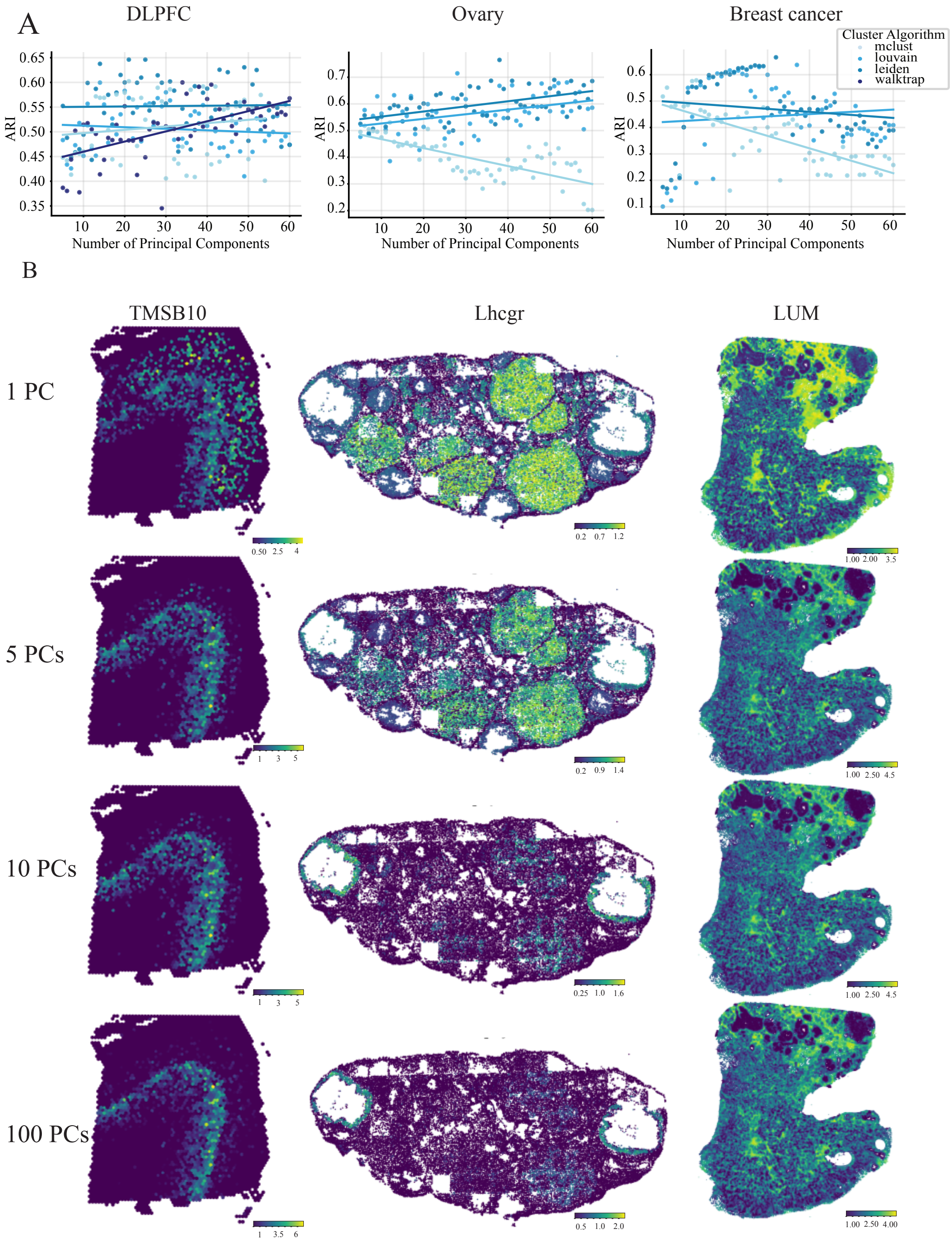

Extended Data Fig. 16: **PC Sensitivity Analysis. A:** ARI values plotted against number of PCs utilized by RASP for the DLPFC (left), Mouse ovary (middle) and Human breast cancer (right) datasets. Colors indicate clustering algorithm. **B:** Reduced rank reconstructed gene signatures for TIMSB10 (left), Lhgr (middle), and LUM (right) at increasing ranks.

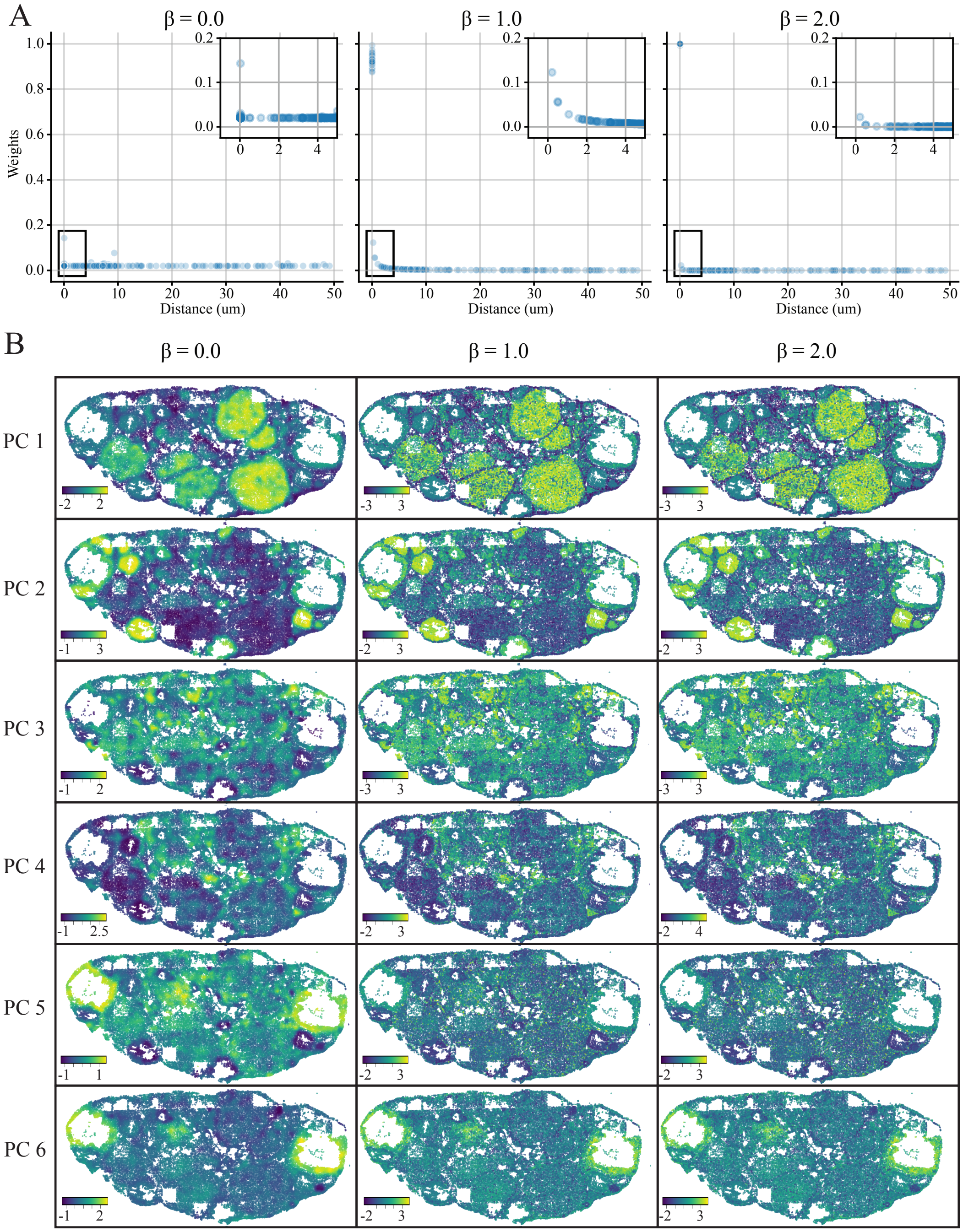

Extended Data Fig. 17: **Effects of inverse weighting on PCs at various $\beta$ values. A:** Inverse weight values plotted against real world distances (um) for $\beta = 0$, $\beta = 1$, and $\beta = 2$. **B:** First six PCs for the mouse ovary dataset visualized spatially on the tissue, smoothed by inverse distances exponentiated by 0 (left), 1 (middle) and 2(right).

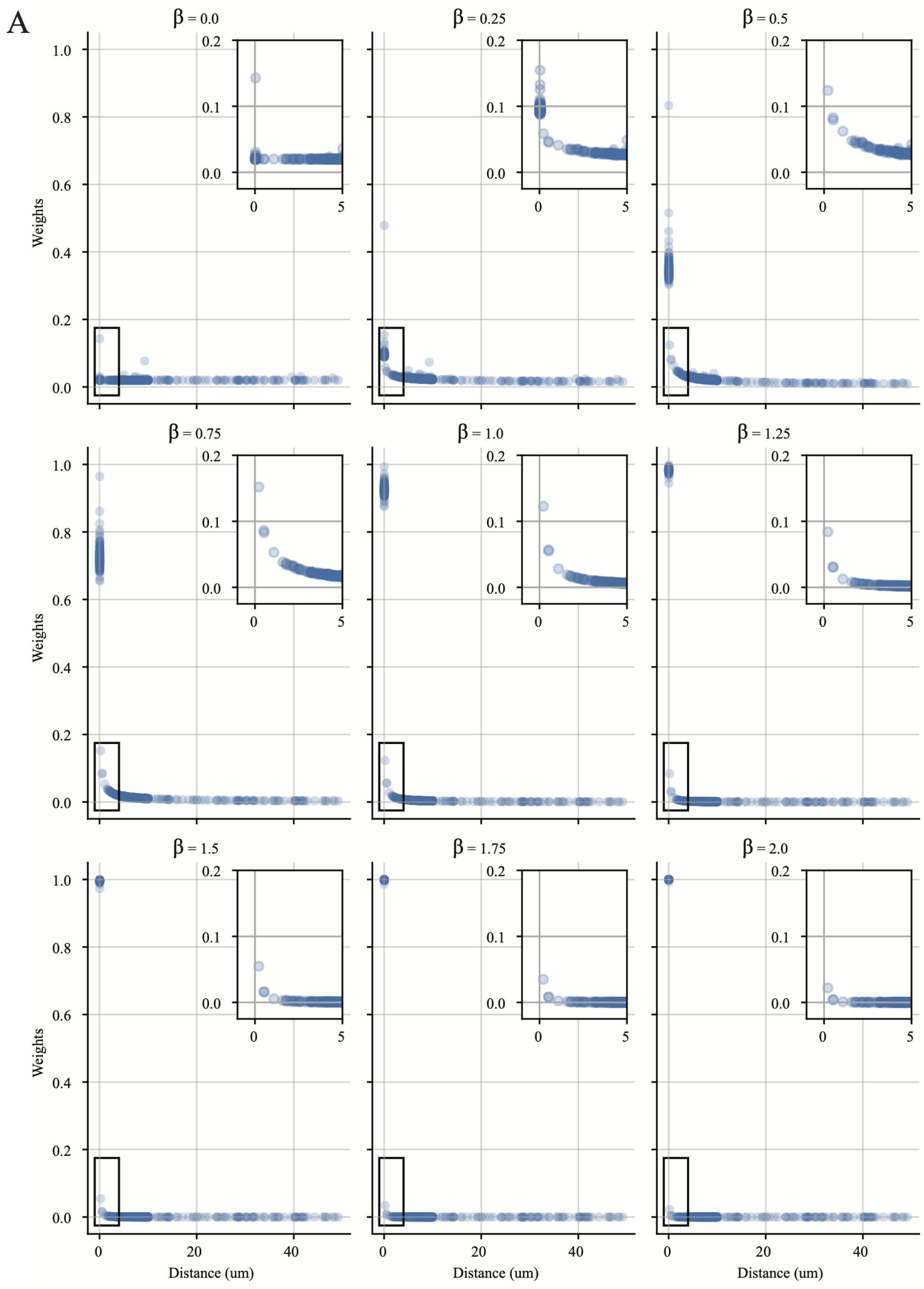

Extended Data Fig. 18: **Inverse weights at different $\beta$ values. A:** Inverse weight values plotted against real world distances (um) at various values of $\beta$

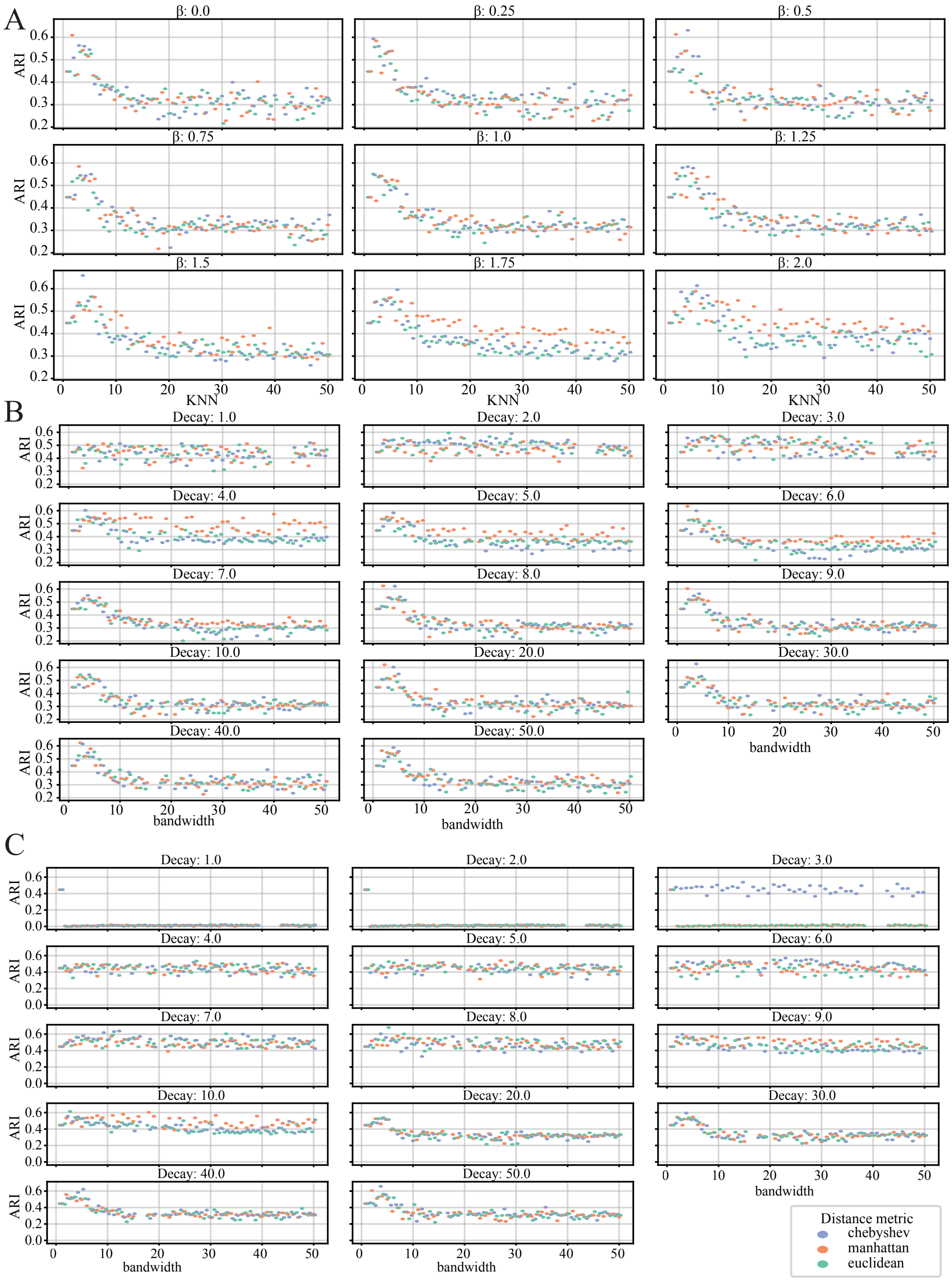

Extended Data Fig. 19: **Differential distance measurements and distance weights applied to DLPFC dataset. A:** Inverse distance weighting across beta parameters **B:** Gaussian kernel weighting across bandwidth and decay parameters, **C:** Quadratic kernel weighting across bandwidth and decay parameters. For all panels colors indicate distance metric used

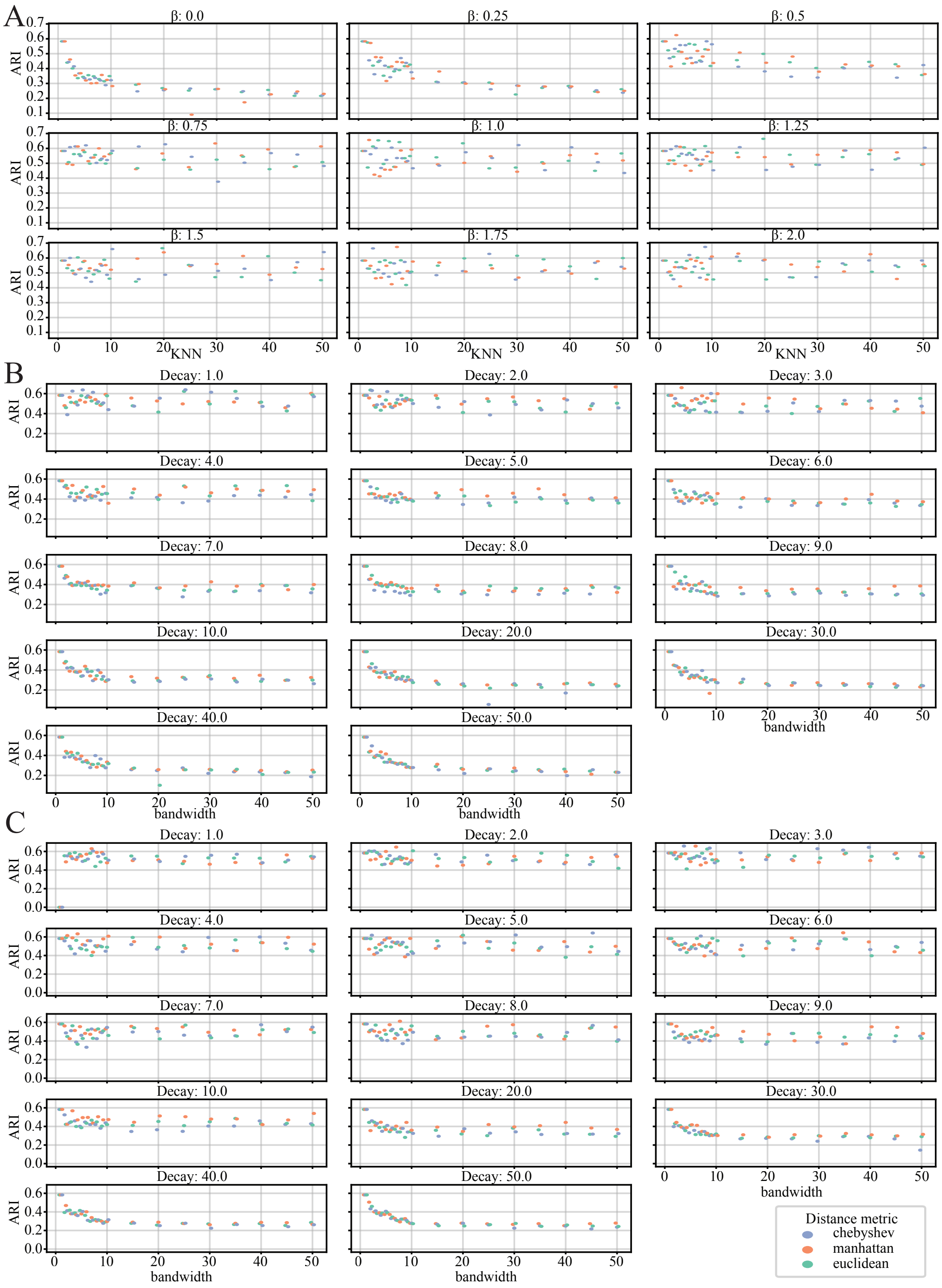

Extended Data Fig. 20: **Differential distance measurements and distance weights applied to the mouse ovary dataset. A:** Inverse distance weighting across beta parameters **B:** Gaussian kernel weighting across bandwidth and decay parameters, **C:** Quadratic kernel weighting across bandwidth and decay parameters. For all panels colors indicate distance metric used

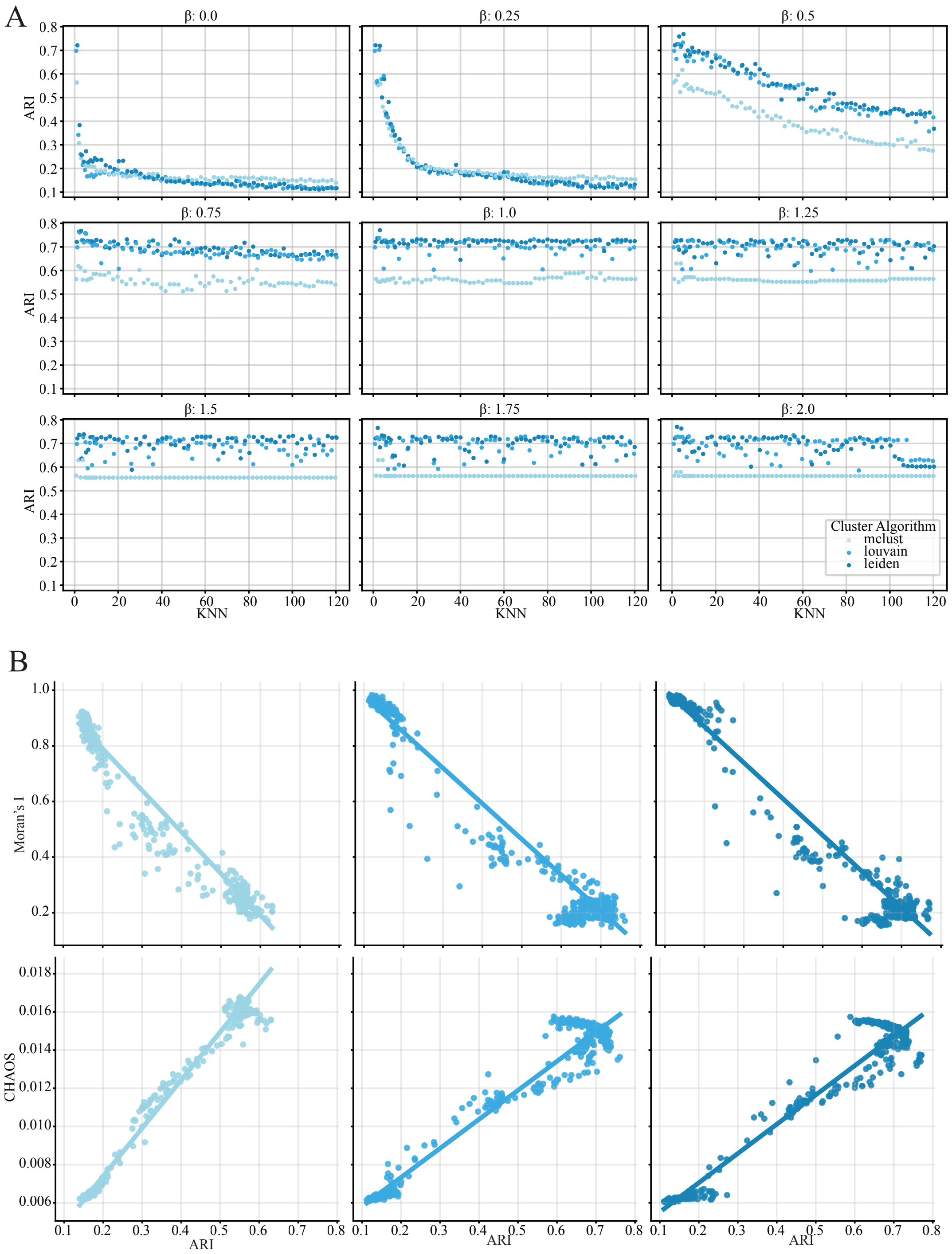

Extended Data Fig. 21: **Sagittal mouse brain supplement 1. A:** ARI values for cell type plotted against kNN distance threshold. Colors indicate clustering algorithm, each subplot corresponds to a distinct $\beta$ value. **B:** Moran's I and CHAOS values plotted against ARI value, colors indicate clustering algorithm. Lines represent robust linear regression best fit.

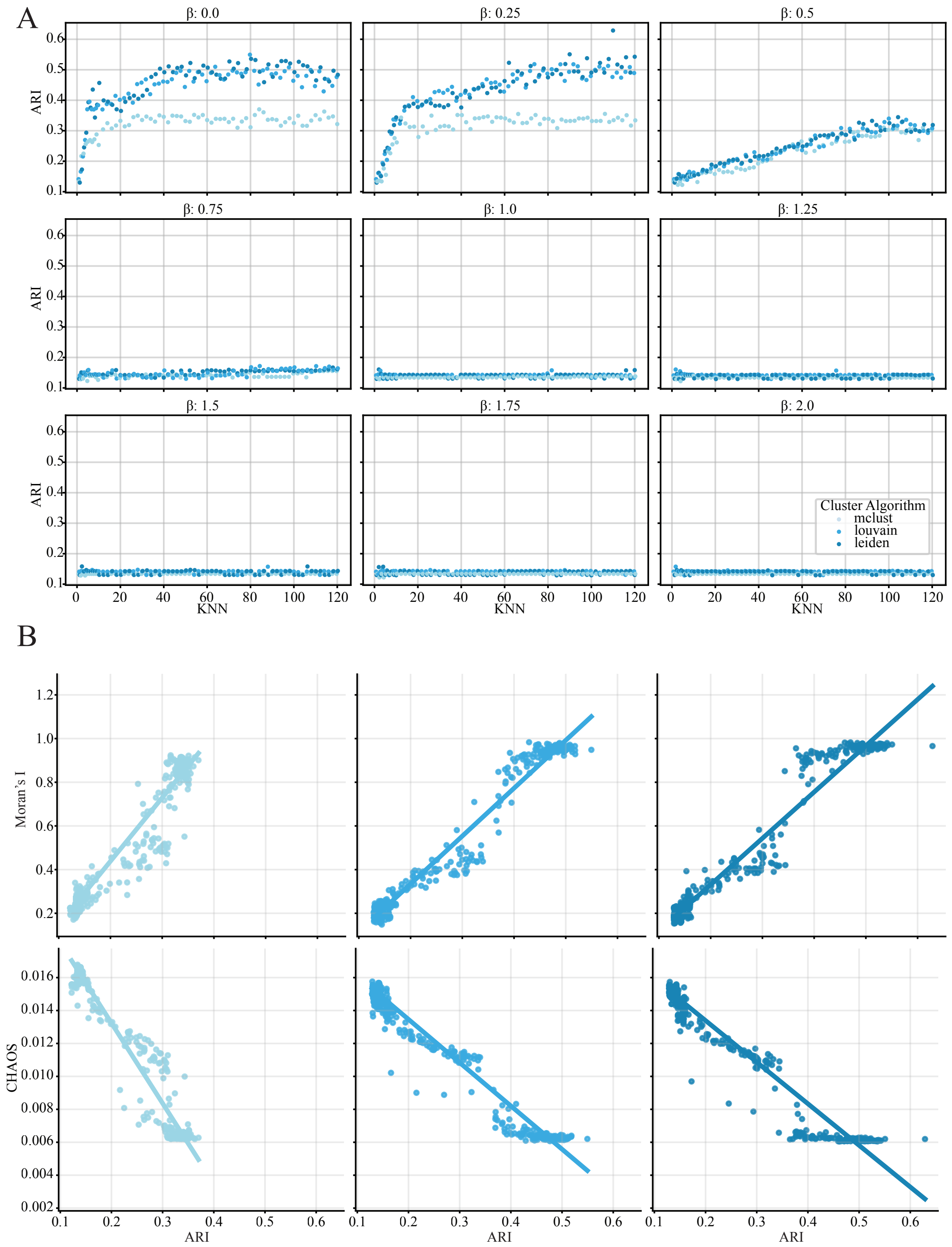

Extended Data Fig. 22: **Sagittal mouse brain supplement 2. A:** ARI values for region annotation plotted against kNN distance threshold. Colors indicate clustering algorithm, each subplot corresponds to a distinct $\beta$ value.**B:** Moran's I and CHAOS values plotted against ARI value, colors indicate clustering algorithm. Lines represent robust linear regression best fit.

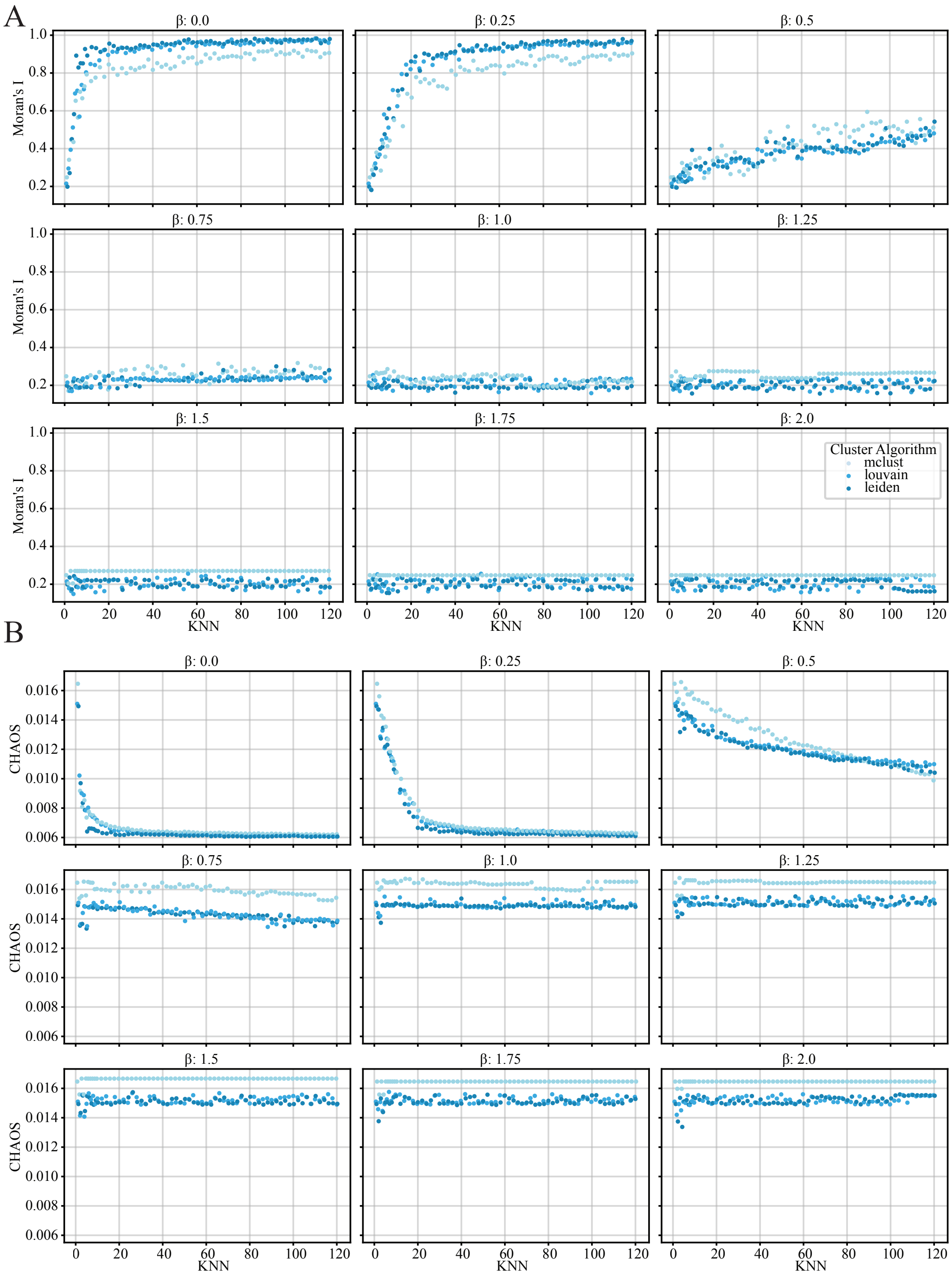

Extended Data Fig. 23: **Sagittal mouse brain supplement 3. A:** Moran's I statistic plotted against kNN distance threshold. Colors indicate clustering algorithm, each subplot corresponds to a distinct $\beta$ value. **B:** CHAOS score plotted against kNN distance threshold. Colors indicate clustering algorithm, each subplot corresponds to a distinct $\beta$ value.

### 2 Supplemental Results

#### 2.1 A note on Moran's I and CHAOS score

Many publications utilize Moran's I and the CHAOS score as measures of cluster quality in the absence of ground truth annotations, where calculating the ARI is not feasible. This study employs these metrics for assessing the olfactory bulb dataset. To this end, we visualized both Moran's I and CHAOS score over varying kNN thresholds from 1 to 50, and for $\beta$ values ranging from 0 to 2 (see Extended Data Fig. 2, 4, 5, and 7).

Across all datasets analyzed, the scores exhibited a similar trend: for small $\beta$ values between 0 and 1, as the kNN threshold increased, the Moran's I value rose logarithmically, while the CHAOS score decayed exponentially. In contrast, for larger $\beta$ values from 1 to 2, the trends persisted but with lower Moran's I values and higher CHAOS scores being observed. Notably, in the case of the ovary dataset, both the CHAOS and Moran's I values remained invariant to changes in the kNN threshold when $\beta$ exceeded 1, with scores in the other datasets trending towards little to no change as well.

Additionally, we visualized the relationship between ARI and Moran's I, as well as between ARI and CHAOS score for all datasets with available ground truth annotations. Interestingly, for the DLPFC and ovary datasets, we found that Moran's I and ARI were negatively correlated (see Extended Data Fig. 1,3). In contrast, the CHAOS score was positively correlated with ARI in the ovary dataset, regardless of the clustering algorithm used. However, in the DLPFC dataset, only the Louvain and Leiden clustering methods demonstrated a positive association with the CHAOS score, while Walktrap and MCLust did not exhibit such a relationship.

These results underscore both the importance and the limited utility of CHAOS score and Moran's I in assessing cluster quality. Moreover, they suggest that the specific clustering task—such as distinguishing between cell types versus spatial domain annotations—and the choice of clustering algorithm will significantly impact the usefulness of these two metrics in evaluating cluster quality.
